# Dynamic regulation of origin firing factors links CDK activity to dormant origin activation

**DOI:** 10.1101/2025.06.10.657920

**Authors:** Md Shahadat Hossain, Courtney G. Sansam, Kimberlie A. Wittig, Tyler D. Noble, Kevin A. Boyd, Christopher L. Sansam

## Abstract

Dormant replication origins help ensure complete genome duplication when replication forks stall, yet how these origins are activated remains poorly understood. Here, we identify a novel regulatory mechanism by which cyclin-dependent kinase (CDK) activity controls the abundance and chromatin recruitment of the origin firing factors TRESLIN and MTBP to promote dormant origin activation. Inhibition of WEE1 kinase during S phase increases CDK activity, which blocks the PCNA-dependent degradation of TRESLIN and enhances its chromatin association along with MTBP. This increased loading is required for elevated helicase recruitment and DNA synthesis under CDK-hyperactive conditions. These effects are reversed by CDK inhibition and depend on both TRESLIN and MTBP. We define a conserved sequence within TRESLIN required for its CDK-sensitive degradation. Significantly, the recruitment of TRESLIN–MTBP and loading of helicase exceed levels observed in unperturbed S phase, supporting a model in which dormant origin firing is actively upregulated through CDK-mediated stabilization of the initiation machinery. These findings uncover a new control point in replication origin usage with implications for genome stability and therapeutic kinase inhibition.

## Introduction

DNA replication initiation through replication origins is a critical process for ensuring accurate genome duplication and maintaining genome stability^1–5^. This process involves a series of coordinated events that begin in the G1 phase of the cell cycle. During this phase, origin licensing occurs, wherein licensing factors, CDC6 and CDT1, facilitate the recruitment of the mini-chromosome maintenance (MCM) helicase complex to replication origins^6,7^. This marks the first step toward replication, as these origins are poised for activation. At the G1/S phase transition, a shift occurs: origin firing factors assemble at these licensed sites, recruiting essential components of the active helicase complex, including CDC45 and the GINS complex^8–10^. Helicase assembly triggers the unwinding of DNA and the initiation of DNA synthesis^11–13^.

Despite the licensing of many origins, it has been estimated that only a fraction—between ∼10-30%—are selected for firing during a normal, unperturbed S phase. The remaining licensed origins, often referred to as "dormant origins," are held in reserve, unused under typical conditions^1,14,15^. These dormant origins function as backups and can be initiated if nearby active replisomes encounter problems during replication, particularly under replicative stress. ^16–18^. This flexibility ensures that the genome can be fully replicated even when replication forks are stalled or compromised. The regulation of dormant origin activation is controlled through inhibitory phosphorylation of the cyclin-dependent kinases during S phase^19,20^.

In unperturbed cells, the rate of replication initiation is not only governed by CDK activity, but also by the availability of origin firing factors. In organisms such as yeast and *Xenopus laevis* embryos, it has been observed that these origin firing factors are present in much lower quantities relative to the number of licensed origins^21,22^. This limiting availability is essential for preventing hyper-replication, a condition that can lead to DNA replication stress and genomic instability^23^. Indeed, studies have shown that overexpression of these limiting factors can result in an excess of replication initiation events, leading to the same detrimental effects^1,24^. In human cells, two key firing factors—TRESLIN (also known as TICRR) and MTBP—are important for origin activation^25–27^. Recent work from our laboratory demonstrated that the chromatin-bound levels of TRESLIN and MTBP decline as cells progress from G1 phase to S phase, and this decrease is caused by proteasome-mediated degradation of TRESLIN, triggered by the E3 ubiquitin ligase CRL4^CDT2^ ^28^. The destruction of TRESLIN at the G1/S transition is paradoxical because TRESLIN-MTBP is generally thought to be necessary for origin firing throughout S phase. However, these unexpected findings suggest that TRESLIN destruction may play a role in limiting the proportion of licensed origins that are activated in S phase.

Here, we used HCT116 cells expressing endogenously tagged TRESLIN, MTBP, or CDC45 to further address the regulation and functional importance of these firing factors. Our results reveal that CDK plays a protective role for TRESLIN, preventing its degradation and ensuring its association with chromatin. Overactivation of CyclinA-CDK2 not only prevents the destruction of TRESLIN but also enhances the chromatin association of TRESLIN and MTBP during S phase. Further, we find that TRESLIN associates with PCNA, and CDK blocks the association to prevent TRESLIN destruction by PCNA-CRL4^CDT2^. Moreover, we identified a specific segment within TRESLIN required for its degradation. Finally, we found that CDK dependent increased recruitment of TRESLIN and MTBP is important for the over-recruitment of CDC45, a component of the replicative helicase which is needed for activation of dormant origins. Collectively, our results provide a deeper mechanistic understanding of how CDK contributes to maintaining proper levels of firing factors and regulates dormant origin firing.

## Results

### WEE1 inhibition of CDK1/2 limits TRESLIN & MTBP recruitment to chromatin during S phase

Our previous study showed that CDK inhibition lowers TRESLIN levels during S phase. This suggested that CDK regulates dormant origin firing by controlling chromatin-bound TRESLIN levels through the CRL4^CDT2^ E3 ligase and the proteasome^28^. To further investigate how CDK influences the levels of TRESLIN and MTBP in S phase, we treated cells with inhibitors that reduce (CDK1/2i; NU-6102^29^) or increase (WEE1i; MK-1775^30^) CDK activity and then measured both total and chromatin-bound TRESLIN or MTBP (Fig. 1). In these experiments, we utilized HCT116 cell lines in which all endogenous TRESLIN or MTBP was tagged with monomeric Clover (mClover)^3^, enabling quantitative measurements of the protein levels using an antibody against GFP^28,31^. We utilized flow cytometry to measure the level of total TRESLIN or MTBP in individual whole cells in addition to the level of TRESLIN or MTBP from the chromatin bound (CB) fraction, which we defined as the detergent resistant pool. We grouped cells into G1 (2N), G2/M (4N), or three distinct phases of S (Early, Mid, or Late) based on the DNA content profile from propidium iodide staining (Supplementary Fig. 1c). Consistent with our prior data^28^, total and CB-TRESLIN levels drop as cells progress from G1 to S phase (Untreated in Fig. 1a-d). During S phase, TRESLIN levels progressively increase, and in G2, they rise to twice their level of G1 (Fig. 1a, b). CB-TRESLIN follows a similar pattern, except its levels remain low in G2 (Fig. 1c, d).

**Fig. 1.**
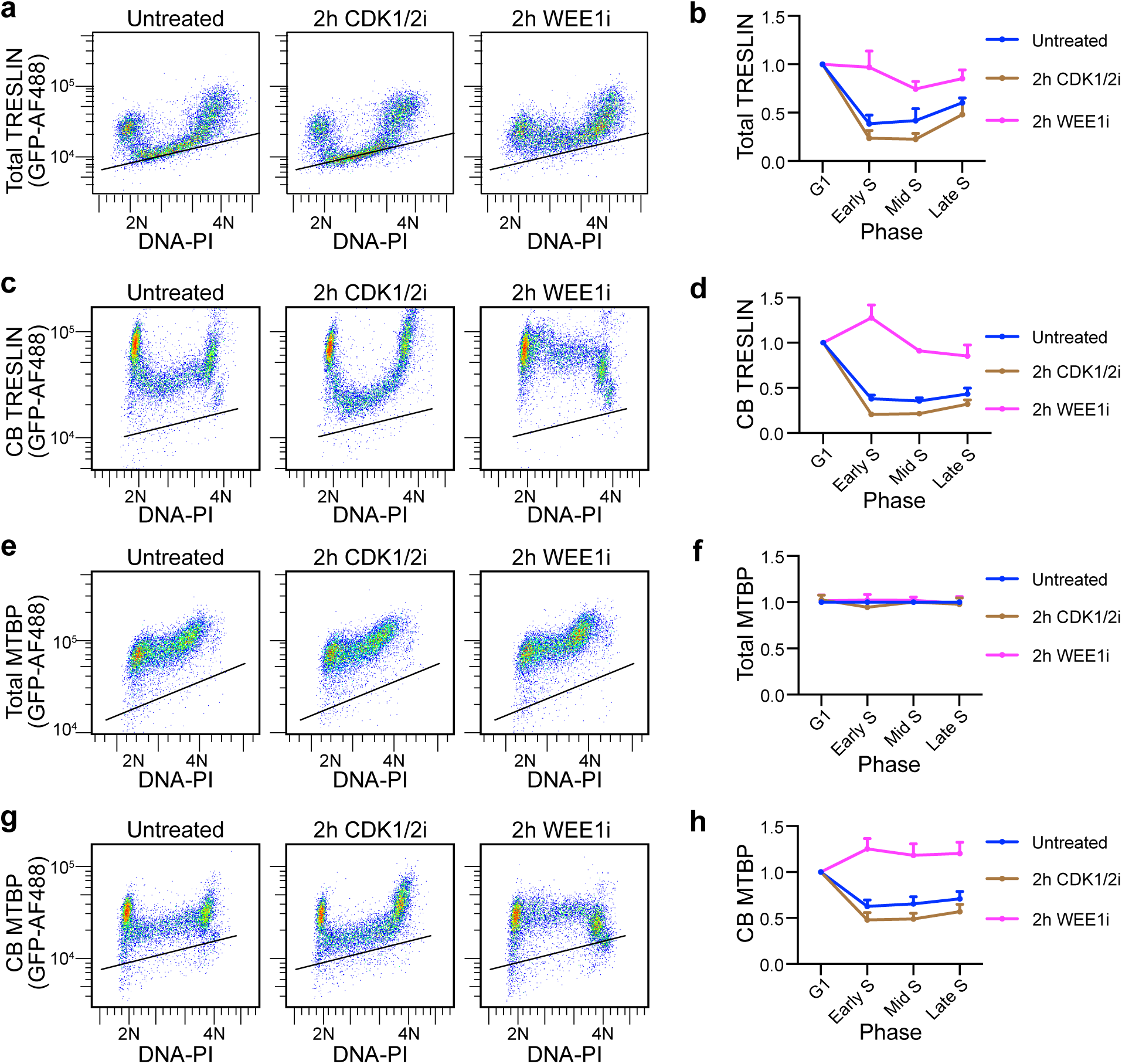
Cell cycle regulation of total and chromatin-bound TRESLIN and MTBP levels in response to CDK1/2 or WEE1 inhibition. (**a, b)** Flow cytometry of total TRESLIN in HCT116 cells with mClover-tagged endogenous TRESLIN. TRESLIN was detected using anti-GFP, and DNA was stained with propidium iodide (PI). **(a)** Pseudocolor plots showing total TRESLIN (y-axis, log scale) vs. DNA content (x-axis, linear scale) after 2-hour treatments as indicated. (b) Quantification of total TRESLIN across the cell cycle. Cells were binned into G1 (2N), early S (>2N), mid S (2N–4N), and late S (<4N). Median TRESLIN signal was background-subtracted, normalized to G1, and plotted as mean ± SD from independent experiments. **(c, d)** Chromatin-bound (CB) TRESLIN measured by flow cytometry. (c) Pseudocolor plots as in (a), showing CB-TRESLIN. (d) Quantification across the cell cycle, as in (b). **(e, f)** Total MTBP levels in mClover-tagged HCT116 cells by flow cytometry, as in (a, b). **(g, h)** CB-MTBP levels, as in (c, d). Line plots (b, d, f, h) represent ≥ 3 independent replicates.

We found that the effects of inhibiting CDK1/2 activity with NU-6102 (hereafter referred to as CDK1/2i) on TRESLIN or MTBP varied across the cell cycle. Two-hour CDK1/2i treatment did not affect TRESLIN in G1 but caused both total and CB-TRESLIN to drop to even lower levels during S phase (Fig. 1a-d). In contrast to its effects on S phase cells, CDK1/2i did not substantially affect total TRESLIN levels in cells with 4N DNA content (Fig. 1a), but it did result in CB-TRESLIN reassociation at that stage (Fig. 1c). In comparison, we found that CDK1/2 inhibition did not affect total MTBP levels (Fig. 1e, f), but it did affect the amount of CB-MTBP (Fig. 1g, h). Like CB-TRESLIN, CB-MTBP levels in G1 were unaffected by CDK1/2i, while its levels in S phase were reduced (Fig. 1g, h). CB-MTBP binding during G2 was increased by CDK1/2i (Fig. 1g). Altogether, we found that reducing CDK activity led to a decrease in the levels of both TRESLIN and MTBP on chromatin during S phase and an increase in their levels on chromatin during G2.

To determine the effects of increasing CDK activity on TRESLIN and MTBP, we used the WEE1 kinase inhibitor MK-1775 (hereafter referred to as WEE1i). WEE1 kinase restrains CDK1/2 activities to ensure faithful DNA replication and mitotic entry^19,32^, and inhibition of WEE1 boosts S phase CDK activity and induces dormant origin firing^33^. If CDK promotes dormant origin firing by increasing available TRESLIN or MTBP, treatment with WEE1i should elevate their levels on chromatin. To test this, we treated the cells with WEE1i for times ranging from 1 to 4 hours and observed increased CB-TRESLIN and CB-MTBP during S phase (Fig. 1c, d and g, h and Supplementary Fig. 1a, b). Interestingly, WEE1i dampened the normal decline in total TRESLIN at the G1/S transition (Fig. 1a), suggesting that WEE1 inhibition prevents the proteasomal degradation of TRESLIN during S phase. The total levels of MTBP were not affected by WEE1i (Fig. 1e, f). These data show that CDK activity levels determine the amount of CB-TRESLIN and CB-MTBP during S phase.

We considered the possibility that the over-recruitment of CB-TRESLIN and CB-MTBP induced by elevated CDK1/2 activity, was caused by changes in origin licensing. To test this, we treated cells with WEE1i and measured the chromatin-bound mini-chromosome maintenance complex subunit MCM7 (CB-MCM7), which marks all potential replication origins during the S phase (Supplementary Fig. 1d, e). WEE1i did not lead to any changes in MCM7 binding during in S phase (Supplementary Fig. 1d, e), consistent with findings by Moiseeva et al.^19^, who demonstrated that WEE1 inhibition does not alter the chromatin loading of MCM2. These results indicate that CDK1/2 promotes TRESLIN and MTBP recruitment to chromatin during S phase without affecting licensing.

### CDK2-Cyclin A2 is required to over-load and stabilize TRESLIN-MTBP on chromatin

Our initial experiments using NU-6102^29^ inhibited both CDK1 and 2 (CDK1/2i; Fig. 1). Given that both CDKs contribute to the G1 to S phase transition, we wanted to determine the relative contributions of each to the levels of CB-TRESLIN and CB-MTBP. Thus, we tested additional inhibitors. Treatment with a selective CDK2/4/6 inhibitor (PF-3600; hereafter referred to as CDK2/4/6i)^34^,which at low concentrations is more specific to CDK2 inhibition^35,36^, reversed the stabilizing effect of WEE1i on CB-TRESLIN and CB-MTBP (Fig. 2a-d).

**Fig. 2.**
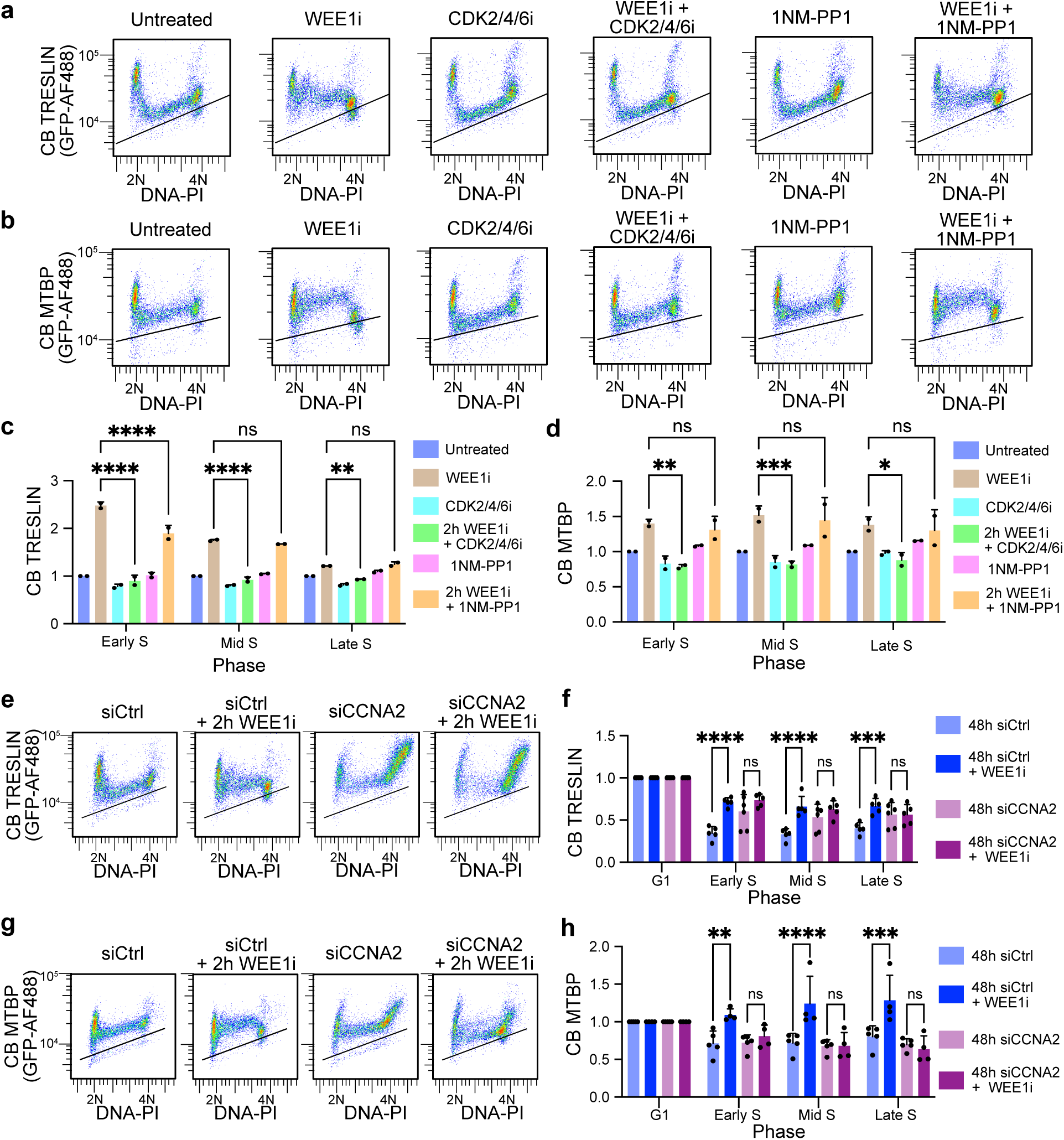
CDK2-Cyclin A2 activity promotes chromatin-bound TRESLIN and MTBP during S phase. **(a, b)** Pseudocolor flow cytometry plots showing chromatin-bound (CB) TRESLIN **(a)** or CB-MTBP **(b)** versus DNA content in HCT116 cell lines with endogenously mClover-tagged TRESLIN or MTBP. CB protein levels were detected using anti-GFP antibody. Cells were treated with the indicated inhibitors. The black line indicates background fluorescence from untagged control cells stained with anti-GFP. **(c, d)** Median CB-TRESLIN **(c)** and CB-MTBP **(d)** levels across cell cycle phases (G1, S, G2/M) from two biological replicates of the data shown in (a, b). Values were background-subtracted and normalized to the untreated condition within each phase. **(e, g)** Pseudocolor flow cytometry plots showing CB-TRESLIN **(e)** or CB-MTBP **(g)** in cells transfected with control (siCtrl) or Cyclin A2 siRNA, with or without WEE1 inhibitor (WEE1i). **(f, h)** Median CB-TRESLIN **(f)** and CB-MTBP **(h)** levels from panels (e) and (g), respectively, across cell cycle phases. Bars represent mean ± SD of replicate medians from >3 independent experiments. In (c, d, f, h), statistical significance was assessed using two-way ANOVA with Tukey’s multiple comparisons test. *P < 0.05; **P < 0.01; ***P < 0.001; ****P < 0.0001.

We initially tested the effect of CDK1 on TRESLIN with the CDK1-selective inhibitor RO-3306 (hereafter referred to as CDK1i), which had effects like CDK1/2i and CDK2/4/6i (Supplementary Fig. 2a-f). However, it has been reported that CDK1i may have off-target effects that affect DNA replication^37,38^. Thus, to more specifically assess the role of CDK1, we generated CDK1-analog sensitive (CDK1^as^) alleles in our TRESLIN and MTBP tagged cell lines by knocking out endogenous CDK1 and introducing a transgene expressing *Xenopus* CDK1^as^ ^39^. In these cell lines, CDK1^as^ can be selectively inhibited by the ATP analog 1NM-PP1^39^ thereby allowing for the specific contributions of CDK1 to the stabilization of TRESLIN and MTBP to be evaluated. Confirming its effectiveness in inactivating CDK1, NM-PP1 treatment of CDK1^as^ cells produced the same effect on mitosis as the CDK1 inhibitor RO-3306 and the opposite effect of the WEE1 inhibitor MK-1775, leading to an accumulation of G2/M phase cells and a reduction in mitotic cells, as detected by anti-MPM-2 (mitotic protein monoclonal 2) staining (Supplementary Fig. 2g–h). Unlike CDK2/4/6i or CDK1i treatment, NM-PP1 treatment of the CDK1^as^ cells did not decrease CB-TRESLIN or CB-MTBP during S phase but did partially reverse the over-loading of TRESLIN onto chromatin during early S phase upon WEE1 inhibition (Fig. 2a-d and Supplementary Fig. 2a-d). These data suggest that when WEE1 is inhibited, CDK1 becomes active and contributes to the over-recruitment and stabilization of CB-TRESLIN during S phase, but in unperturbed cells, TRESLIN and MTBP binding to chromatin in early S phase is primarily controlled by CDK2.

and CB-MTBP, we next addressed whether Cyclin A2 was responsible for the over-recruitment of TRESLIN and MTBP by using siRNA-mediated knockdown (Supplementary Fig. 2i). Cyclin A2 knockdown alone did not affect the levels of CB-TRESLIN and CB-MTBP; however, in combination with WEE1 kinase inhibition the Cyclin A2 knockdown reduced the over-recruitment of TRESLIN and MTBP in S phase (Fig. 2 e-h). These results indicate that Cyclin A2 is required for the over-recruitment of CB-TRESLIN and CB-MTBP during S phase.

### Mutual dependency of TRESLIN and MTBP for CDK-driven chromatin loading

Genetic and biochemical experiments in human cells and *X. laevis* extracts suggest that TRESLIN/MTBP function as an obligate heterodimer ^26,40,41^. Nonetheless, it is unknown whether TRESLIN or MTBP can function independently, and our recent findings indicate that CB-MTBP is TRESLIN-dependent during the G1 phase but not during S phase^31^. As CDK1/2 overactivation stimulates the recruitment of TRESLIN and MTBP to chromatin during S phase (Fig. 1c, d and g, h), we investigated whether they rely on each other for the heightened chromatin association in response to increased CDK activity. Using siRNA knockdown of TRESLIN or MTBP, alongside WEE1 kinase inhibition, we assessed whether knockdown of either protein prevented over-recruitment of the other (Fig. 3 and Supplementary Fig. 3). siRNA knockdown efficacy was demonstrated by flow cytometry, and the extent of knockdown was not affected by WEE1 inhibition (Supplementary Fig. 3a-h).

**Fig. 3.**
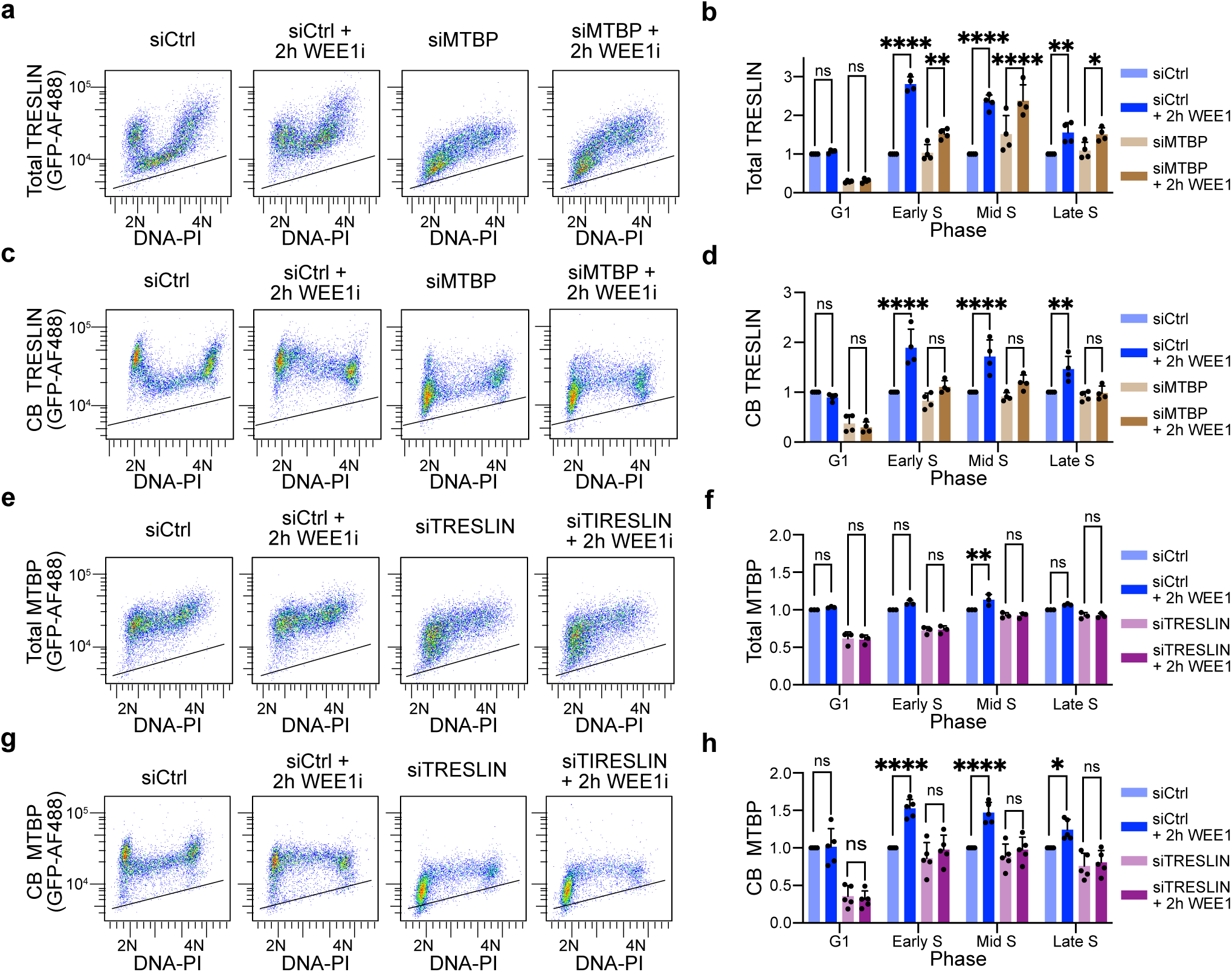
Mutual Dependency of TRESLIN and MTBP for over-recruitment onto chromatin. **(a, b)** Flow cytometry of total TRESLIN levels in HCT116 cells with endogenously mClover-tagged TRESLIN after MTBP knockdown and/or WEE1 inhibition. **(a)** Pseudocolor flow cytometry plots showing total TRESLIN (anti-GFP, y-axis, log scale) versus DNA content (PI, x-axis, linear scale) in cells transfected with siControl or siMTBP, with or without WEE1i (MK1775, 2 h). Black lines indicate background fluorescence from untagged control cells stained with anti-GFP. **(b)** Median total TRESLIN levels across the cell cycle. Cells were binned into G1 (2N), early S (>2N), mid S (2N–4N), and late S (<4N). Values were background-subtracted and normalized to the siControl condition within each phase. **(c, d)** CB-TRESLIN levels measured by flow cytometry after MTBP knockdown and/or WEE1 inhibition. **(c)** Pseudocolor plots as in (a), showing CB-TRESLIN. **(d)** Median CB-TRESLIN levels across the cell cycle, as in (b). **(e, f)** Flow cytometry of total MTBP levels in HCT116 cells with endogenously mClover-tagged MTBP transfected with siControl or siTRESLIN, with or without WEE1i. **(e)** Pseudocolor flow cytometry plots of total MTBP signal. **(f)** Median total MTBP levels across the cell cycle, as in (b). **(g, h)** CB-MTBP levels in TRESLIN knockdown cells, as in (c, d). Bars in (b, d, f, h) represent mean ± SD of replicate medians from ≥3 independent experiments. Statistical analysis was performed using two-way ANOVA with Tukey’s multiple comparisons test. *P < 0.05; **P < 0.01; ***P < 0.001; ****P < 0.0001.

Total MTBP and TRESLIN levels are not interdependent during S phase, indicating they do not rely on each other for stability. Although MTBP knockdown reduced TRESLIN levels in G1, it had no effect during S phase (Fig. 3a, b). WEE1i treatment increased total TRESLIN levels throughout S phase, even in the absence of MTBP, indicating that MTBP is not required for CDK-dependent stabilization of TRESLIN (Fig. 3a, b). Likewise, TRESLIN knockdown did not alter total MTBP levels during S phase (Fig. 3e, f), suggesting that MTBP stability is also maintained independently of TRESLIN.

We next tested whether MTBP is required for the chromatin binding of TRESLIN. MTBP knockdown blocked the WEE1i-induced increase in chromatin-bound TRESLIN (Fig. 3c, d). Similarly, TRESLIN knockdown prevented the increase in chromatin-bound MTBP following WEE1 inhibition (Fig. 3g, h). Overall, these results show that while MTBP is not necessary for stabilizing TRESLIN during WEE1 inhibition, TRESLIN and MTBP rely on each other for their increased recruitment to chromatin during S phase when CDK activity increases.

### PCNA colocalizes with TRESLIN and is required for TRESLIN removal from chromatin during S phase

We recently demonstrated that PCNA knockdown impedes TRESLIN degradation during S phase^28^. Consistent with this, siRNA-mediated depletion of PCNA also prevented the loss of chromatin-bound (CB) TRESLIN between the G1 and S phases (Fig. 4a, b). While these findings suggest a role for PCNA in regulating TRESLIN, it remains unclear whether PCNA directly promotes TRESLIN removal from chromatin or targets it for degradation. To explore this, we examined whether TRESLIN and PCNA co-localize at replication foci.

**Fig. 4.**
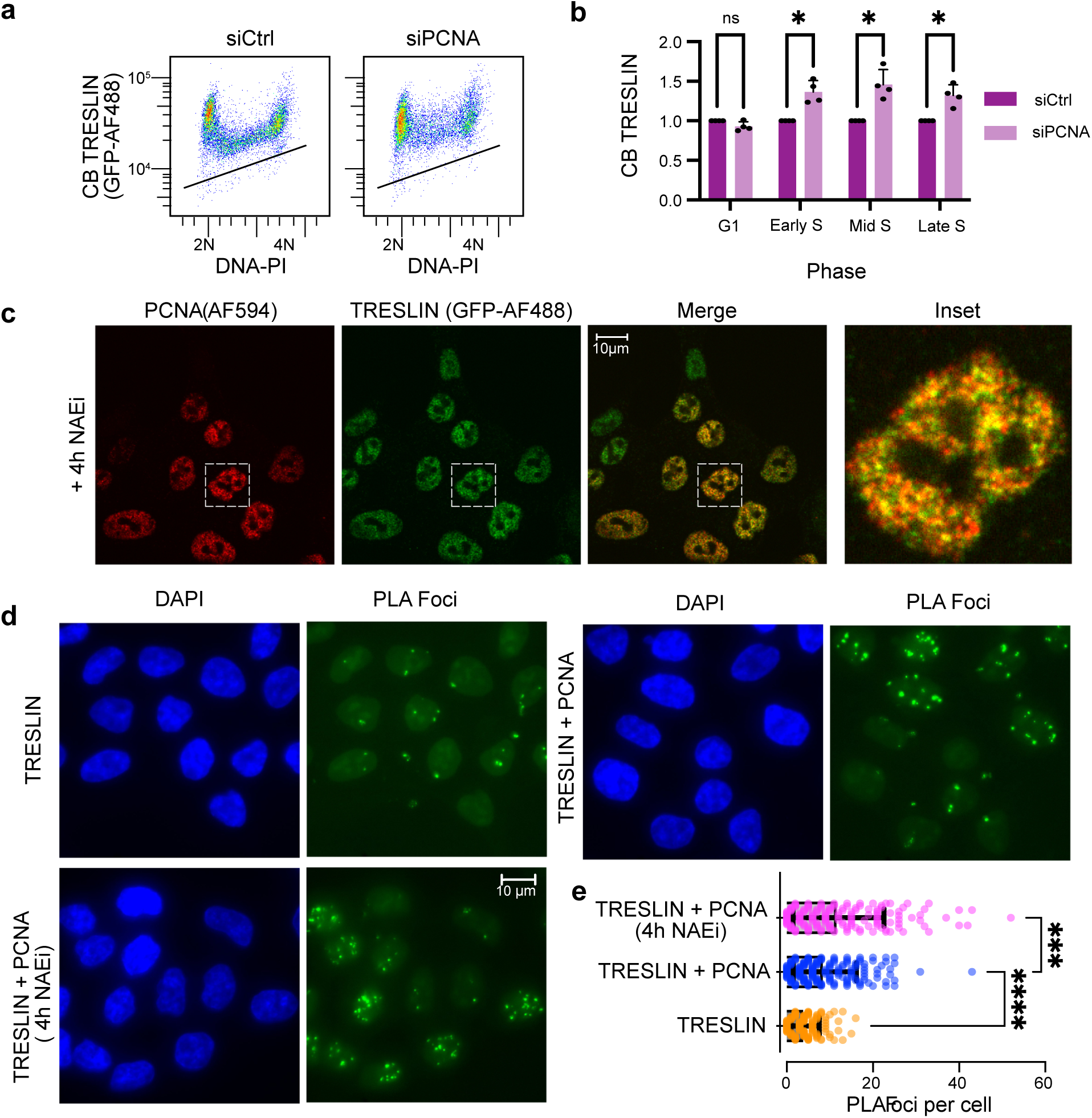
PCNA colocalizes with TRESLIN and promotes its removal from chromatin during S phase. **(a)** Pseudocolor flow cytometry plots of chromatin-bound (CB) TRESLIN versus DNA content (PI) in HCT116 cells expressing mClover-tagged TRESLIN. Cells were transfected with control (siControl) PCNA (siPCNA) siRNA. Black lines indicate background anti-GFP signal measured in untagged control cells processed in parallel. **(b)** CB-TRESLIN quantification from (a) across four replicates. Data points represent the background-subtracted median anti-GFP signal in individual DNA content gates corresponding to G1, early S, mid S, and late S. Values are normalized to the median siControl signal within each phase. Bars represent the mean + SD of replicate median quantification. **(c)** Immunofluorescence images showing nuclear localization of TRESLIN-mClover (green, anti-GFP) and PCNA (red) in HCT116 cells treated with the neddylation inhibitor MLN4924 (NAEi) for 4 hours. Inset shows a representative nucleus. **(d)** Proximity ligation assay (PLA) for TRESLIN-PCNA interaction with the following antibody combinations: anti-GFP alone (TRESLIN, n = 157), anti-GFP plus anti-PCNA (TRESLIN + PCNA, n = 194), and anti-GFP plus anti-PCNA following NAEi treatment (n=173). **(e)** Quantification of PLA from (d). Each point represents an individual nucleus; the number of PLA foci per cell is plotted. Bars represent mean + SD. Statistical analysis was determined using one-way ANOVA followed by Tukey’s multiple comparisons test. *P<0.05, **P<0.01, ***P<0.001, ****P<0.0001.

Because TRESLIN is degraded during S phase, it is difficult to detect by immunofluorescence at this stage. To overcome this, we used the neddylation inhibitor MLN4924 (hereafter referred to as NAEi) to block Cullin-RING ligase–mediated degradation and stabilize TRESLIN protein^28^. Immunofluorescence using antibodies against GFP (to detect endogenously tagged TRESLIN-mCLOVER) and PCNA revealed nuclear co-localization of the two proteins (Fig. 4c). To more rigorously assess their spatial proximity, we performed proximity ligation assays (PLA), which allow sensitive and specific detection of protein interactions within 40 nm in situ^42^. PLA using GFP and PCNA antibodies produced a nuclear signal (Fig. 4d TRESLIN + PCNA) which was reduced in control reactions using the GFP antibody alone (Fig. 4d TRESLIN). Quantification of nuclear foci confirmed the PLA signal increase (Fig. 4e). The PLA signal was further enhanced by NAEi treatment—consistent with TRESLIN stabilization and increased association with PCNA (Fig. 4d, e).

### CDK protects TRESLIN via blocking TRESLIN and PCNA association

We next investigated the mechanism by which CDK activity protects TRESLIN from degradation. TRESLIN’s association with PCNA suggests it may be targeted for destruction by being brought within proximity of PCNA-associated CRL4^CDT2^, like known substrates such as CDT1, p21, and SET8^43^. As shown above for TRESLIN (Fig. 1), CDK activity also protects CDT1, p21, and SET8 from degradation^44^. In these cases, CDK1-mediated phosphorylation of the CRL4 adapter protein CDT2 reduces its affinity for PCNA, thereby limiting CRL4^CDT2^ activity as CDK1 activity rises in late S and G2 phases^44,45^.

Consistent with this model and previously published data, flow cytometry showed that CDT1 levels drop sharply at the G1/S transition and reaccumulate in G2/M (Supplementary Fig. 4b). WEE1 inhibition (WEE1i) further elevated CDT1 levels in G2/M (Supplementary Fig. 4b, c). However, unlike the WEE1i effects on TRESLIN (Fig. 1a,b), WEE1i treatment has less effect on CDT1 levels in S phase (Supplementary Fig. 4b), likely due to parallel CDT1 degradation pathways (e.g., SCF^46^) or differential regulation by CDK.

Because CDK-mediated phosphorylation of CDT2 reduces its PCNA binding, we hypothesized that CDT2 overexpression might overcome CDK-mediated protection of TRESLIN and CDT1. To test this, we generated a transgenic HCT116 cell line expressing endogenously tagged TRESLIN-mClover and harboring a PiggyBac-based tetracycline-inducible CDT2 expression cassette (Supplementary Fig. 4a). Upon induction, CDT2 overexpression reversed the WEE1i-induced increase in CDT1 during G2/M but did not block TRESLIN accumulation during S phase (Supplementary Fig. 4b, c, d). This suggested that additional mechanisms contribute to TRESLIN stabilization in S phase.

We hypothesized that CDK protects TRESLIN from destruction by limiting its localization with PCNA. To test this hypothesis, we quantified colocalization of TRESLIN and PCNA in S phase cells treated with CDK, neddylation, or WEE1 inhibitors. Endogenous TRESLIN-mClover and PCNA were visualized by immunofluorescence using anti-GFP and anti-PCNA antibodies (Fig. 5). Analyses were restricted to S phase cells identified by PCNA foci. Colocalization was quantified using Pearson’s correlation coefficient (Fig. 5b) and Mander’s coefficients M1 (PCNA overlapping TRESLIN; Fig. 5c) and M2 (TRESLIN overlapping PCNA; Fig. 5d).

**Fig. 5.**
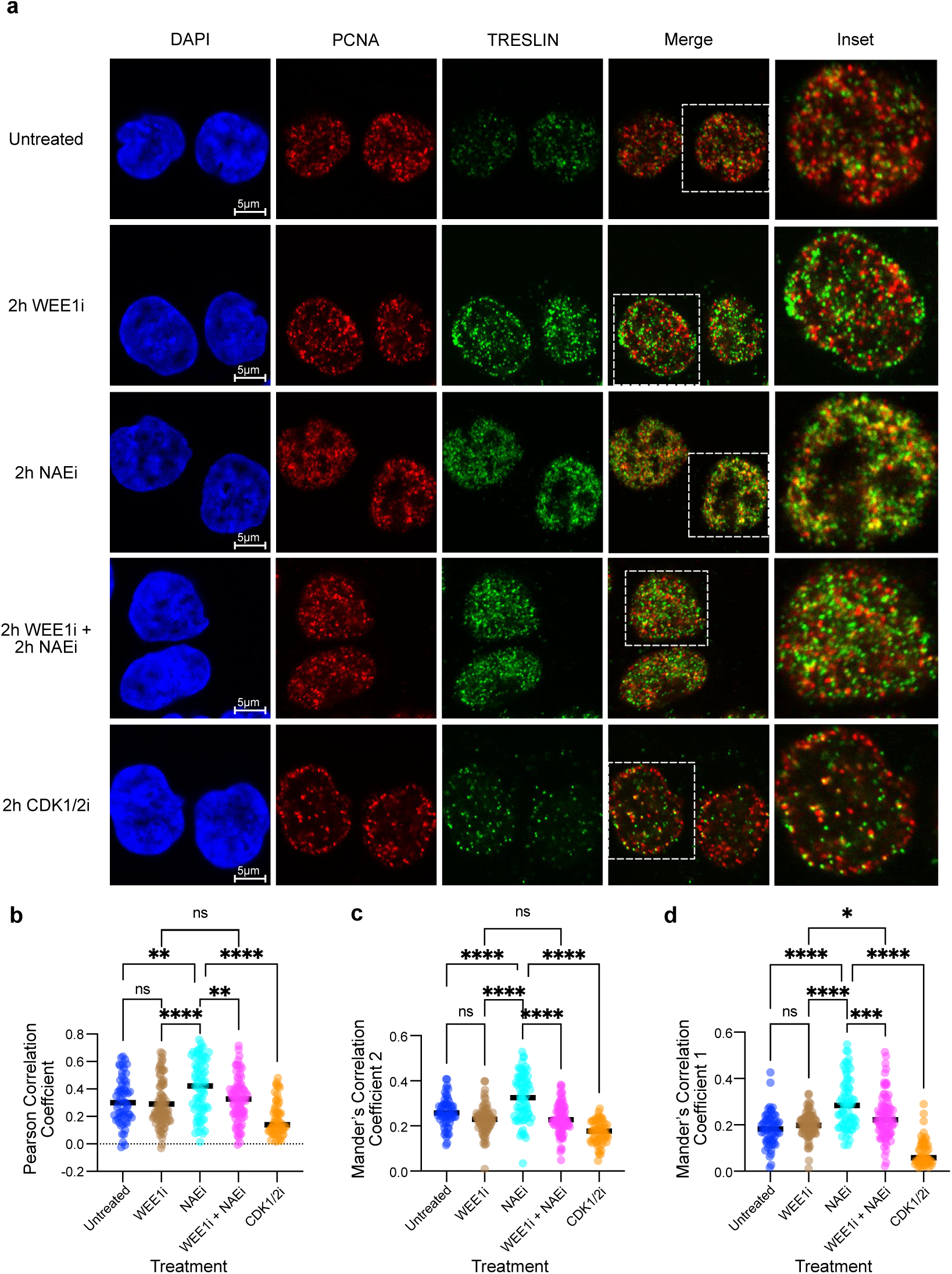
CDK protects TRESLIN by preventing its localization with PCNA at replication foci. **(a)** Representative confocal images of HCT116 cells with endogenously tagged TRESLIN-mClover after 2-hour treatment with WEE1 inhibitor (WEE1i; MK1775), neddylation inhibitor (NAEi; MLN4924), CDK1/2 inhibitor (CDK1/2i; NU6102), or their combinations as indicated. Cells were immunostained with anti-GFP (detecting TRESLIN-mClover) and anti-PCNA antibodies. Insets show magnified views of single nuclei (dashed boxes). Images are restricted to S phase cells identified by the presence of PCNA foci. **(b–d)** Quantification of TRESLIN–PCNA colocalization in S phase cells under the indicated treatments. **(b)** Pearson’s correlation coefficient. **(c)** Mander’s M2 coefficient (fraction of TRESLIN overlapping with PCNA). **(d)** Mander’s M1 coefficient (fraction of PCNA overlapping with TRESLIN). Each point represents a single nucleus; horizontal lines indicate medians. Statistical comparisons were performed by one-way ANOVA with Tukey’s post hoc test.

As expected, TRESLIN levels were low in untreated S phase cells, and its overlap with PCNA was minimal (Fig. 5a top row; 5b-d blue). CDK1/2 inhibition further reduced TRESLIN levels and its colocalization with PCNA (Fig. 5a bottom; 5b-d orange), consistent with increased destruction of TRESLIN. Conversely, neddylation inhibition (NAEi) increased both TRESLIN levels and its colocalization with PCNA (Fig. 5a third row), as reflected by elevated Pearson and Mander’s coefficients (Fig. 5b-d turquois), suggesting accumulation of TRESLIN at replication foci normally cleared by CRL4^CDT2^-mediated degradation. In contrast, WEE1i increased TRESLIN levels but did not enhance colocalization with PCNA (Fig. 5a second row; 5b-d brown), suggesting that WEE1i stabilizes TRESLIN by preventing its recruitment to PCNA-associated foci.

If CDK protects TRESLIN by reducing its PCNA association, then WEE1i should also suppress the colocalization of TRESLIN stabilized by NAEi. Indeed, combined treatment with WEE1i and NAEi reduced colocalization to levels observed with WEE1i or untreated controls (Fig. 5a-d). These results indicate that NAEi and WEE1i stabilize TRESLIN through distinct mechanisms: NAEi blocks CRL4^CDT2^ activity but allows the localization of TRESLIN to PCNA foci, while WEE1i prevents TRESLIN recruitment to replication foci, thereby shielding it from CRL4^CDT2^-mediated ubiquitination.

### A ΔSBI TRESLIN mutant is resistant to destruction in S phase

To further understand the mechanisms and functions of CDK-regulated TRESLIN destruction, we next focused on identifying sequences in TRESLIN necessary for its degradation. We first tested whether TRESLIN degradation depends on sequences within N- or C-terminal regions. To investigate this, we generated N- and C-terminal truncation mutants tagged with mfGFP and stably expressed them in 293 Flp-In T-Rex cells (Fig. 6a). Protein at the expected molecular weights was confirmed by capillary electrophoresis (Fig. 6b), and overall expression levels were quantified using live-cell GFP fluorescence via flow cytometry (Fig. 6c).

**Fig. 6.**
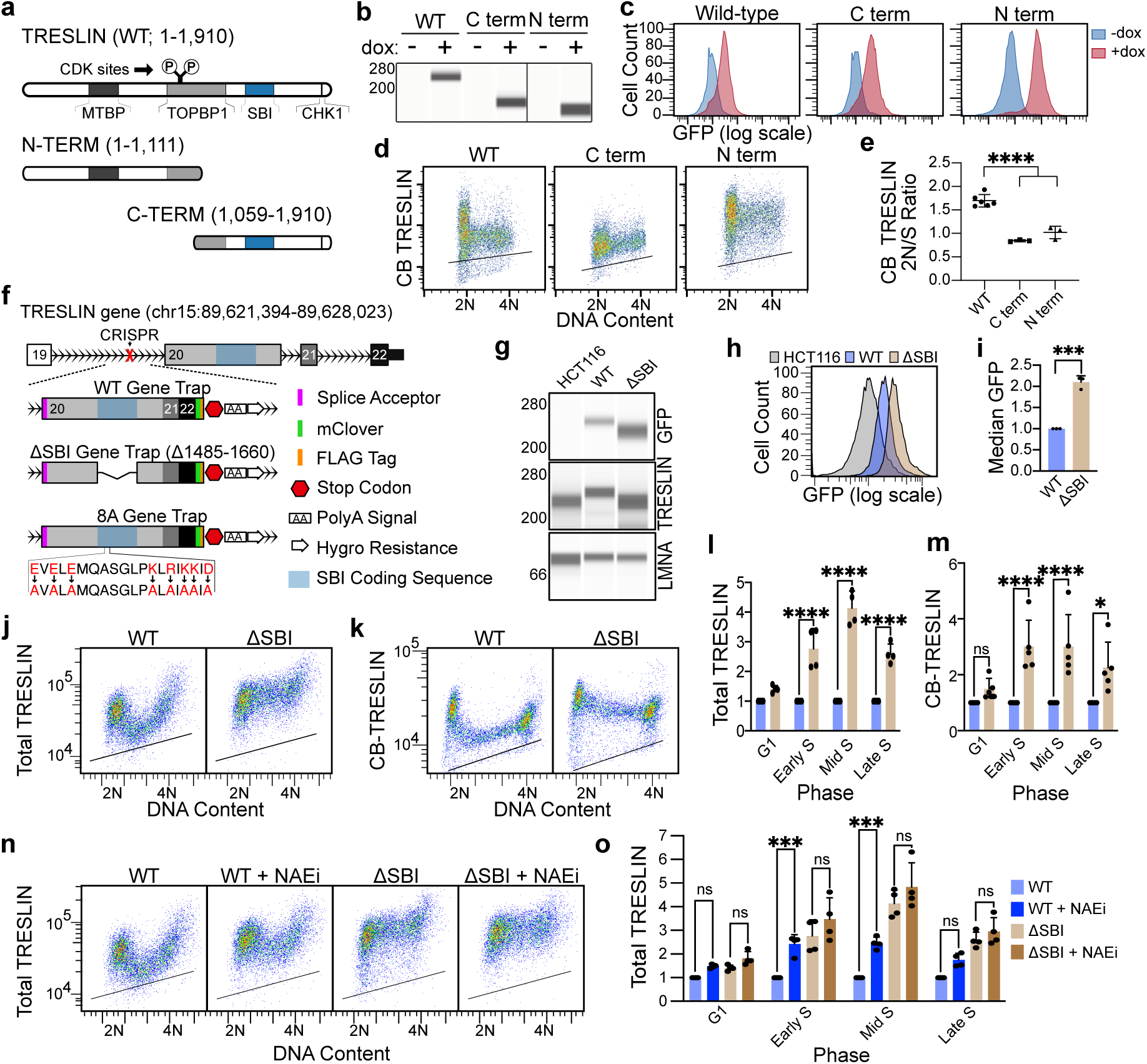
Deletion of the SBI region stabilizes TRESLIN during S phase by blocking its degradation. **(a–e)** Analysis of GFP-tagged TRESLIN fragments in HEK293 Flp-In T-REx cells. **(a)** Schematic of TRESLIN domains, including interaction regions for MTBP, TOPBP1, BET proteins (SBI: aa 1485–1661), and CHK1. Tet-On constructs express wild type or N- or C-terminal halves of TRESLIN with N-terminal GFP. **(b)** Capillary electrophoresis (Jess) of whole-cell lysates ± doxycycline (Dox) with anti-GFP. **(c)** Flow cytometry of GFP fluorescence in live cells from (b); histograms show log-scale GFP intensity ± Dox. **(d)** Flow cytometry of CB GFP-TRESLIN fragments in cells from (a); pseudocolor plots show anti-GFP signal (y-axis, log scale) versus DNA content (x-axis, linear scale). Black lines indicate background from –Dox cells. **(e)** G1/S ratio of CB GFP-TRESLIN signal from (d); values represent background-subtracted median anti-GFP signal in G1 (2N) and S phase (2N–4N) gates across replicates. **(f–m)** Analysis of wild-type and ΔSBI TRESLIN in HCT116 knock-in cell lines. **(f)** Strategy for generating knock-in lines expressing wild-type, ΔSBI, or 8A TRESLIN-mClover. A CRISPR-induced intronic cut enabled insertion of a splice acceptor and modified coding sequences. **(g)** Capillary electrophoresis of whole-cell lysates from lines in (f), probed with anti-GFP, anti-TRESLIN, and anti-LMNA. **(h)** Live GFP fluorescence measured by flow cytometry. **(i)** Median GFP quantification from (h); n = 2. (j, k) Flow cytometry of total **(j)** and CB **(k)** TRESLIN in wild-type and ΔSBI lines. Pseudocolor flow cytometry plots show anti-GFP (log scale) versus DNA content (linear scale); black lines indicate background from untagged control lines. **(l, m)** Quantification of total **(l)** and CB **(m)** TRESLIN from (j, k). Median anti-GFP signal in G1, early S, mid S, and late S was background-subtracted and normalized to wild-type for each gate. **(n, o)** TRESLIN expression in wild-type and ΔSBI lines after 2 h treatment with neddylation inhibitor (NAEi; MLN4924). **(n)** Flow cytometry of total TRESLIN (anti-GFP). **(o)** Quantification of (n), as in (l). Statistical analysis was performed using two-way ANOVA with Tukey’s multiple comparisons test. *P < 0.05; **P < 0.01; ***P < 0.001; ****P < 0.0001.

Although all constructs were integrated at the same Flp-In locus, the N-terminal half of TRESLIN was expressed at higher levels than the full-length or the C-terminal half of TRESLIN (Fig. 6c). As expected, cell cycle profiling of its abundance by flow cytometry showed that full-length TRESLIN expression declined at the G1/S transition (Fig. 6d; WT). In contrast, both truncation mutants maintained relatively constant expression across the cell cycle (Fig. 6d C term and N term; quantified by 2N/S ratio in Fig. 6e). Although both truncation mutants remained stable from G1 to S phase, their baseline expression levels differed (Fig. 6d). The N-terminal truncation, which lacks the MTBP-binding domain, showed low expression in both phases. Conversely, the C-terminal truncation exhibited high expression in G1 and remained elevated in S phase. These findings suggest that the C-terminal half of TRESLIN contains sequences essential for its proteolytic degradation.

While analyzing BET protein interactions with TRESLIN, we serendipitously identified a stabilized TRESLIN mutant lacking a portion of its C-terminal domain. We previously defined amino acids 1485–1661 as necessary and sufficient for BET protein binding—the SBI region^47^. Using a gene trap/CRISPR knock-in strategy, we deleted the SBI region and inserted an mClover tag at the endogenous *TRESLIN* locus in HCT116 cells (Fig. 6f). PCR confirmed successful insertion of the gene trap into both *TRESLIN* alleles, and a wild-type control line was generated using the same method (Fig. 6f). Capillary electrophoresis using TRESLIN and GFP antibodies showed that in the gene trap line, the majority of TRESLIN protein carried the mClover fusion tag (Fig. 6g). The ΔSBI mutant displayed a lower molecular weight, consistent with loss of the SBI sequence (Fig. 6g). Notably, ΔSBI TRESLIN was more abundant than wild-type, as confirmed by electrophoresis and GFP flow cytometry (Fig. 6g–h).

Unexpectedly, chromatin-bound ΔSBI TRESLIN levels remained high during S phase, indicating that loss of the SBI region blocks normal S phase–dependent degradation (Fig. 6j-m). We examined the protein’s response to neddylation inhibition to test whether the ΔSBI mutation affects the CRL4^CDT2^ S phase pathway responsible for TRESLIN degradation. As previously shown^28^, treatment with NAEi stabilized wild-type TRESLIN during S phase, preventing its normal G1/S decline (Fig. 6n). However, the ΔSBI mutant was already elevated in S phase and showed no further significant increase upon NAEi treatment (Fig. 6n). These data demonstrate that deletion of the SBI region renders TRESLIN resistant to CRL4^CDT2^-mediated degradation.

To determine whether the ΔSBI mutation stabilizes TRESLIN by disrupting its interaction with BET proteins, we mutated eight conserved charged residues in the SBI region (TRESLIN-8A), previously shown to be required for BRD4 binding (Fig. 6f)^47^. CRISPR knock-in of this 8A mutant into HCT116 cells revealed that it was degraded similarly to wild-type TRESLIN during S phase and responded normally to CDK1/2 modulation (Supplementary Fig. 5c–g). These results indicate that the sequence required for TRESLIN degradation is distinct from the BRD4-interacting residues within the SBI region. Altogether, our data show that the SBI deletion disrupts a sequence in TRESLIN needed for its destruction at the G1/S transition and that the specific sequence within the SBI region is distinct from that required for the BET protein interaction.

### TRESLIN stabilization is insufficient for hyper-replication

Given that TRESLIN levels were stabilized by WEE1i, we investigated whether TRESLIN stabilization was necessary or sufficient to stimulate DNA synthesis. Since we were able to isolate homozygous ΔSBI mutants, this mutation does not severely impact the proliferation of HCT116 cells. A comparison of cell cycle profiles revealed significant, reproducible differences in the proportion of cells in G1 and G2/M phases between the ΔSBI and wild-type (WT) gene trap lines (Fig. 7a). Specifically, the ΔSBI line had a significantly lower percentage of cells in G1 (16.4% vs. 26.3%; Šídák’s p = 0.0131) and a significantly higher percentage of cells in G2/M (25.3% vs 18.5%; Šídák’s p < 0.0001; Fig. 7a). Despite these differences, the ΔSBI mutant cells did not have a difference in median EdU incorporation measured by flow cytometry (Fig. 7b). Furthermore, WEE1i treatment increased EdU incorporation in mid and late S phase in the ΔSBI cells to a similar level as WT cells (Fig. 7b, c). These results show that stabilization of TRESLIN by the ΔSBI mutation does not stimulate DNA replication, nor does it prevent the activation of dormant origins by WEE1i treatment. These data suggest that in HCT116 cells, stabilization of TRESLIN by CDK is not sufficient to activate dormant origins.

**Fig. 7.**
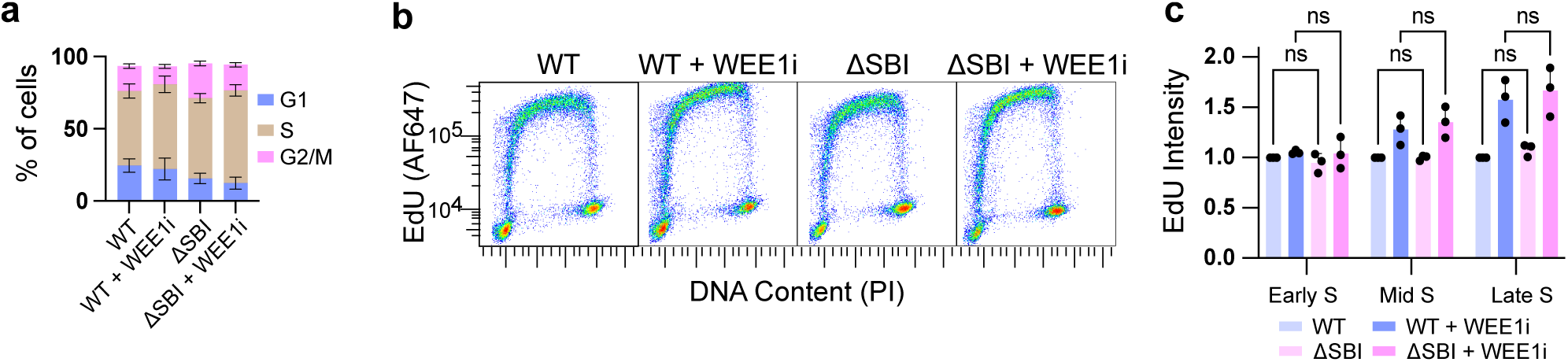
Stabilization of TRESLIN by ΔSBI mutation does not promote DNA replication or prevent WEE1i-dependent origin activation. **(a)** Cell cycle distribution of wild-type (WT) and ΔSBI TRESLIN-mClover HCT116 knock-in cells, measured by flow cytometry using DNA content and EdU incorporation. Stacked bar plots show the mean percentage of cells in G1, S, and G2/M phases across replicates; error bars represent standard deviation. Significant differences in G1 and G2/M proportions were detected by Šídák’s multiple comparisons test. **(b)** Representative pseudocolored flow cytometry plots of EdU incorporation (y-axis, log scale) versus DNA content (x-axis, linear scale; PI staining) in WT and ΔSBI cells, with or without 2-hour WEE1 inhibitor (WEE1i; MK1775) treatment. **(c)** Quantification of EdU signal from (b). Cells were gated into early, mid, and late S phase based on DNA content. Median EdU signal in each bin was background-subtracted and normalized to the WT signal for the corresponding cell cycle phase. Bars show mean + SD from biological replicates. Statistical analysis was determined using two-way ANOVA followed by Tukey’s multiple comparisons test.

### Increased recruitment of TRESLIN and MTBP is essential for dormant origin activation

Next, we tested whether the increased recruitment of TRESLIN or MTBP to chromatin is necessary for the activation of dormant origins. Pulse labeling of cells with EdU confirmed that WEE1 kinase inhibition increases DNA synthesis, while inhibition of CDK2/4/6 reduces it (Fig. 8a). Since CDC45 is loaded onto chromatin during helicase assembly, its chromatin-associated levels can be used as a proxy for fork numbers. Therefore, we generated a CDC45-mClover knock-in HCT116 cell line, enabling us to measure CDC45 on chromatin by flow cytometry using anti-GFP (Supplementary Fig. 6a-c). As expected, WEE1i and CDK2/4/6i inhibitors increased and decreased the levels of CDC45 on chromatin, respectively, paralleling their effects on EdU incorporation (Fig. 8b).

**Fig. 8.**
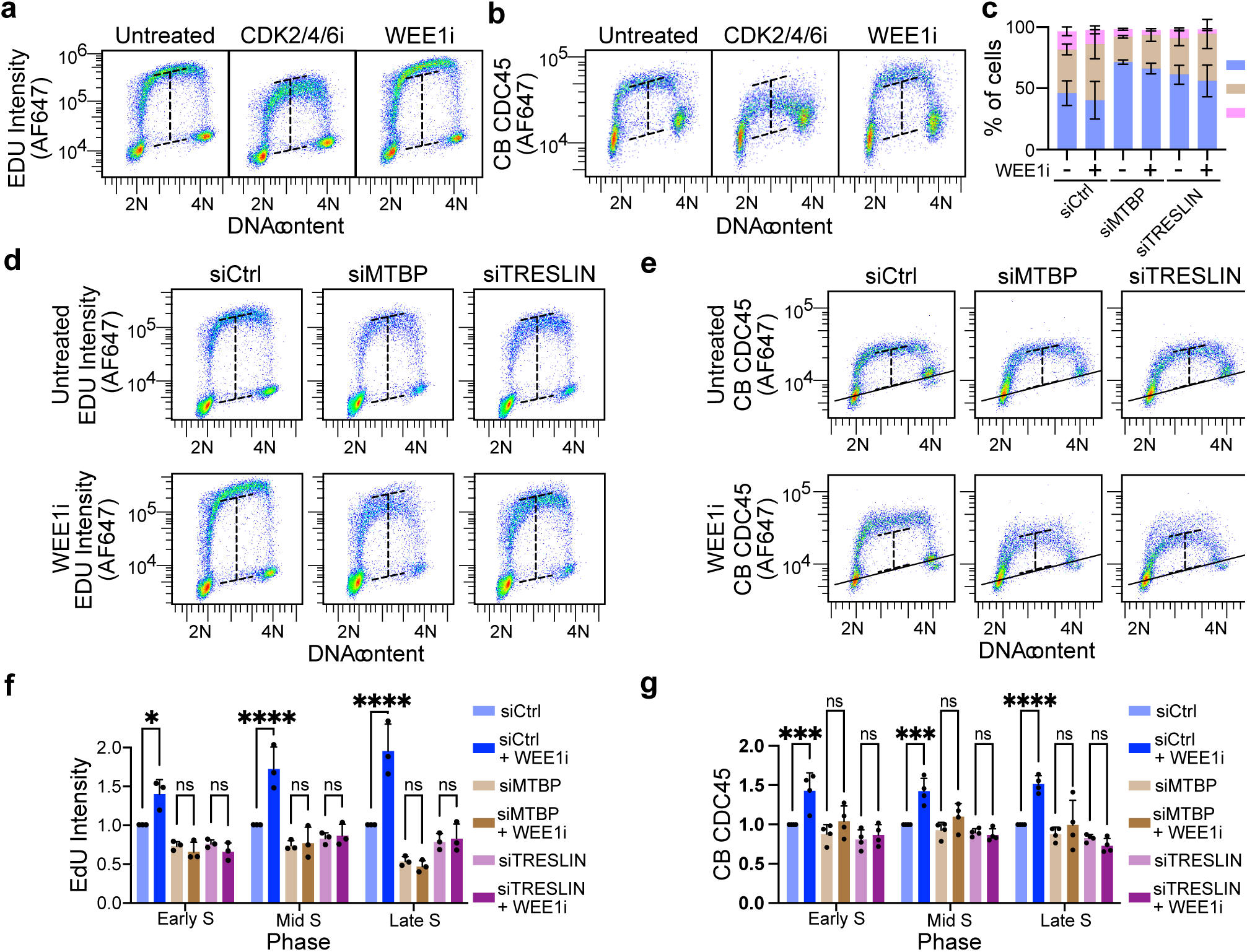
TRESLIN and MTBP are required for CDK-dependent activation of dormant replication origins. **(a)** Flow cytometry pseudocolor dot plots of EdU (log10 scale) vs DNA content (Propidium iodide; linear scale) in untreated HCT116 cells or treated with CDK2/4/6 inhibitor (CDK2/4/6i), or WEE1 inhibitor (WEE1i). Dashed lines indicating the EdU-negative baseline and the mid S-phase EdU peak in the Untreated sample and are shown in all panels to facilitate comparison. **(b)** Chromatin-bound CDC45 (CB-CDC45) levels under the same drug conditions as in a, measured using an anti-GFP antibody in a CDC45-mClover knock-in cell line. Dashed lines mark the CDC45-negative baseline and the peak CDC45 signal in the Untreated sample. The solid black line represents background signal measured in parallel with untagged cell line. **(c)** Stacked bar plots showing the proportion of cells in G1, S, or G2/M phases from four biological replicates of HCT116 cells transfected with siControl, siTRESLIN, or siMTBP. Cell cycle phase was determined from flow cytometry of EdU versus DNA content, as in d. Bars represent means and error bars indicate standard deviation (SD) across replicates. **(d)** Representative pseudocolor dot plots of EdU versus DNA content in siControl, siTRESLIN, or siMTBP cells with or without WEE1i treatment. Dashed lines in all plots indicate EdU levels in the siControl sample. **(e)** Representative pseudocolor dot plots of CB-CDC45 versus DNA content under the same conditions as in d. Dashed lines mark CDC45 levels in the siControl sample. The solid black line marks the background signal measured in untagged HCT116 cells analyzed in parallel. **(f)** Quantification of EdU incorporation from four biological replicates of the experiment shown in d. Median EdU signal was calculated for cells gated in early, mid, and late S phase based on DNA content. Values were background subtracted and normalized to siControl within each S phase fraction. **(g)** Quantification of CB-CDC45 signal from four biological replicates of the experiment shown in e, analyzed as in f. Bars represent means and error bars represent SD across biological replicates. Statistical comparisons were performed using two-way ANOVA with Tukey’s post-hoc test.

To investigate the role of TRESLIN and MTBP in dormant origin activation, we used siRNAs to knock down each protein. By measuring protein levels in the tagged cell lines via flow cytometry, we assessed knockdown efficiency across the cell cycle (Supplementary Fig. 3). Although TRESLIN and MTBP levels were decreased in S phase following siRNA knockdown, they did not go below background levels (Supplementary Fig. 3). siRNA knockdown of either TRESLIN or MTBP resulted in an increased proportion of HCT116 cells in G1 and a reduction in EdU incorporation during S phase (Fig. 8c, d). However, substantial EdU incorporation remained, with TRESLIN knockdown cells maintaining 96%, 81%, and 68% of siControl EdU incorporation levels in early, mid, and late S phase, respectively (Fig. 8d, f). Similarly, MTBP knockdown cells exhibited 87%, 68%, and 64% of siControl levels across early, mid, and late S phase (Fig. 8d, f). Correspondingly, CDC45 levels were reduced in TRESLIN knockdown cells to 80%, 89%, and 82% of siControl levels in early, mid, and late S phase, and to 88%, 93%, and 88% in siMTBP cells (Fig. 8d, g).

The partial reduction in EdU incorporation observed in TRESLIN or MTBP knockdown cells suggests that the proteins become limiting for origin firing during normal S phase progression upon knockdown with siRNA. We then tested whether WEE1 inhibition (WEE1i) could increase the expression or chromatin binding of TRESLIN and MTBP under these conditions. Neither the expression nor chromatin binding of TRESLIN or MTBP was enhanced by WEE1i treatment in knockdown cells (Supplementary Fig. 3). This outcome provided the opportunity to test whether WEE1i could stimulate dormant origin firing without concomitantly increasing TRESLIN or MTBP levels. Although EdU incorporation and CDC45 chromatin binding were elevated in siControl-transfected cells upon WEE1i treatment, no significant increase was observed in cells transfected with siTRESLIN or siMTBP (Fig. 8d-g). We confirmed these results in human retinal pigment epithelial (RPE-1) cells (Supplementary Fig. 7). These results indicate that, in the absence of sufficient TRESLIN and MTBP, CDK cannot effectively increase origin firing, thus supporting the role for CDK-mediated stabilization of TRESLIN in activating dormant origins.

## Discussion

Previous studies in yeast and higher eukaryotes have established that the levels of origin-firing factors, including Sld3/TRESLIN and Sld7/MTBP, determine fork initiation rates and S phase duration in different biological contexts^21,48^. These studies demonstrated that low expression levels of these factors limit dormant origin activation and prolong S phase, underscoring the importance of their abundance in replication origin firing regulation. Given their limited availability, these factors serve as critical nodes for signaling pathways that control replication initiation rates and S phase progression. Our study identifies a novel regulatory mechanism by which CDK2-Cyclin A controls the stability and chromatin association of TRESLIN and MTBP during S phase. Specifically, our data indicates that overactivated CDK2-Cyclin A blocks the association between TRESLIN and PCNA, preventing its degradation and promoting the chromatin association of MTBP. This mechanism regulates CDC45 recruitment to chromatin, thereby influencing dormant origin firing. By linking CDK activity to the stabilization and function of TRESLIN and MTBP, we provide mechanistic insight into how signaling pathways can rapidly modulate these factors to regulate origin firing rates and S phase length.

Our study further clarifies how TRESLIN regulation affects dormant origin firing. Our previous work showed that expression of the phosphomimetic "TESE" TRESLIN mutant, which mimics CDK phosphorylation at key sites that promote its binding to TOPBP1, enhanced origin firing, accelerated replication timing, and shortened S phase in U2OS cells. In contrast, overexpression of wild-type TRESLIN did not stimulate DNA replication in these cells, revealing that TRESLIN phosphorylation was rate-limiting for origin firing when TRESLIN was overexpressed. Interestingly, in HeLa cells, overexpressing wild-type TRESLIN was sufficient to promote origin firing, highlighting cell type-specific regulatory differences^33^. Here, we tested the effect of a stabilized TRESLIN mutant (ΔSBI) in HCT116 cells at endogenous levels and found that it alone was insufficient to increase origin firing. However, high TRESLIN levels were required for dormant origin activation in WEE1-inhibited cells, indicating that TRESLIN stabilization is necessary for dormant origin firing. Overall, our results demonstrate that CDK promotes dormant origin firing through two distinct mechanisms: TRESLIN phosphorylation, which promotes its interaction with TOPBP1, and TRESLIN stabilization by preventing CRL4^CDT2^ mediated ubiquitination and proteasomal degradation. The fact that TRESLIN overexpression in HeLa cells, but not U2OS or HCT116 cells, indicates that the relative contribution of these mechanisms in regulating origin firing appears to vary by cell type.

Our findings are consistent with those of Charrasse et al., who demonstrated that knockdown of the PP2A-B55 inhibitor ENSA led to decreased TRESLIN levels, reduced origin firing, and slower S phase progression due to increased TRESLIN degradation^49^. Since PP2A-B55 dephosphorylates CDK substrates, ENSA knockdown effectively mimics CDK inhibition. Our study expands upon these findings in two ways. First, we demonstrate that TRESLIN degradation occurs primarily during S phase and is linked to DNA replication through CRL4^CDT2^ and PCNA. Second, we show that CDK inhibits TRESLIN degradation during S phase. Together, these findings highlight that TRESLIN levels are not set before S phase entry but are dynamically regulated throughout S phase by pathways that modulate S phase CDK activity and CDK substrate phosphatases such as PP2A-B55. This dynamic regulation underscores the interplay between CDK activity and phosphatase function in controlling origin firing and S phase progression.

While our data clearly show that CDK stabilizes TRESLIN, the precise mechanism by which it does so remains unclear. Other CRL4^CDT2^ substrates, such as CDT1, CDKN1A, and SET8, are ubiquitinated upon recruitment to PCNA via PIP degrons. Although TRESLIN lacks a clear PIP degron, our study provides evidence that TRESLIN and PCNA are closely positioned in replication foci. Additionally, while CDK1 phosphorylation of CDT2 protects other CRL4^CDT2^ substrates from degradation by inhibiting its interaction with PCNA, our data demonstrates that WEE1 inhibition both stabilizes TRESLIN and reduces its colocalization with PCNA, suggesting that CDK2 protects TRESLIN by inhibiting its localization to PCNA foci, which would conceivably reduce its ubiquitination by CRL4^CDT2^. Although Charrasse et al. showed that the TESE phosphomimetic mutant is stabilized, our previously published work indicated that the TESE mutant is degraded at the G1/S transition like wild-type TRESLIN. Therefore, we propose that the CDK phosphorylated residues (T969 and S1001) critical for replication initiation are less significant for regulating TRESLIN stability. Given that CDK phosphorylates other TRESLIN residues, CDK-dependent stabilization may be caused by TRESLIN phosphorylation.

We show here that the chromatin association of both TRESLIN and MTBP is also regulated by CDK. Notably, we also show that CDK regulates the chromatin association of MTBP without affecting its overall levels, and this regulation appears to be independent of TRESLIN stabilization. Furthermore, TRESLIN and MTBP expression levels are highest in G2 when CDK levels are high, yet they are not bound to chromatin at that stage. Additional studies are needed to elucidate the precise mechanisms by which CDK protects TRESLIN from destruction and regulates the chromatin association of MTBP and TRESLIN in specific cell cycle phases.

The data presented here enhances our understanding of how signaling pathways regulate S phase progression. Our findings clarify how developmental signals, replication stress, and oncogenes can influence origin firing rates through regulating the stability of TRESLIN and the chromatin association of TRESLIN and MTBP. This research also provides insights into how inhibitors of kinases such as WEE1, CHK1, and ATR, which are currently under clinical investigation, promote the hyper-activation of dormant origins.

## Methods

### Cell culture

HCT-116 colon cancer cells and hTERT-RPE-1 immortalized human retinal pigment epithelial cells were obtained from ATCC (CCL-247 and CRL-4000) and cultured in McCoy’s 5A or Dulbecco’s Modified Eagle’s F12 Media (Corning), respectively, supplemented with 10% fetal bovine serum (FBS, BioWest). Flp-In T-REx 293 cells (Invitrogen, R78007) were cultured in DMEM containing 10% FBS. Where indicated, cells were treated with MLN2924 (3uM, Caymen), NU-6102 (20uM, Caymen), MK-1775 (1uM, Selleck), RO3306 (10uM, Caymen), PF-3600 (25nM, Caymen), 1nM-PP1 (5uM, Calbiochem), Doxycycline (2.5ug/ml Enzo), Hygromycin (200ug/ml, Enzo).

### Stable cell line generation

The following parental cell lines were used for genetic modification: HCT116 (RRID:CVCL_0291), HCT116 MTBP-mClover (RRID:CVCL_C7SR), HCT116 TICRR-mClover (RRID:CVCL_C7SS), and Flp-In T-REx 293 (RRID:CVCL_U427). Genetic engineering strategies included CRISPR/Cas9-mediated mutation or knock-in, PiggyBac or Sleeping Beauty transposition, and Flp-In recombination. Key lines generated for this study include a CDC45-mClover knock-in, CDK1 analog-sensitive variants of MTBP-mClover and TICRR-mClover, a doxycycline-inducible CDT2 transgenic variant of TICRR-mClover, intronic gene trap HCT116 lines expressing wild-type or mutant (ΔSBI and 8A) TRESLIN isoforms, and inducible wild type or truncation TRESLIN in Flp-In T-REx 293 cells. Detailed cloning strategies, selection methods, and validation procedures are described in the Supplemental Methods.

### siRNA transfections

All siRNAs were transfected using RNAiMax (ThermoFisher Scientific) according to the manufacturer’s instructions. At 24hrs post transfection, the cells were split and incubated for an additional 24hrs. Drugs or inhibitors were added during the final 2 hours prior to harvesting. Cyclin A2 (S2512, Life Technologies), siTreslin (Kumagai et al, 2010, Life Technologies)^27^, siMTBP (M-013953–01-0010; Dharmacon), siPCNA (VC30004, Sigma); siCTRL (12935110, Life Technologies). siRNA sequences are provided in Supplemental Methods.

### Flow cytometry

Live cell, total, and chromatin-bound immuno-flow cytometry experiments were performed as described in Wittig et al.^28^. Primary Antibodies used for flow cytometry include GFP (used for Treslin, MTBP, CDC45, Rockland 600–401-215S), PCNA (PC10, Santa Cruz Sc-56), CDT1 (Abcam ab202067), MCM7 (Santa-Cruz sc-56324), MPM-2 (DAKO M3514). All flow cytometry data was analyzed using FlowJo (TreeStar, Inc.). Gating strategies described by Wittig et al.^28^ were applied in FlowJo to identify single cells, measure protein expression, and distinguish cell cycle phases. Median fluorescence intensity for each phase was calculated.

### EdU incorporation assay

20 µM EDU (Life Technologies) was added to the culture media and incubated during the final 30 minutes of each experiment. The cells were then harvested, washed in 1X PBS and resuspended in a hypotonic lysis buffer [0.1% Sodium Citrate and 0.03% NP-40] for 6 minutes on ice. The resulting nuclei were resuspended in 200ul of a Click-it cocktail containing PBS, 2mM CuSO4, 2uM A647 Azide (Life Technologies), and 50mM ascorbic acid, incubated in the dark at room temperature for 90 minutes, washed and resuspended in a DNA staining solution containing 10ug/ml propdium iodide and 100ug/ml RNAse A prior to running on the analyzer (Cytoflex, Beckman-Coulter). Data was analyzed using FlowJo.

### Microscopy

For GFP and PCNA staining, cells were grown on coverslips. The coverslips were incubated on ice in a pre-permeabilization buffer (0.5% Triton X-100, 20 mM HEPES at pH 7.9, 50 mM NaCl, 3 mM MgCl2, 300 mM sucrose) then fixed in 2% PFA for 20 min and in ice-cold methanol for 10 min, blocked with PBS containing 3% BSA prior to antibody labeling. Primary antibodies (GFP 1:500 Rockland, PCNA 1:500 Santa Cruz). DNA was stained with DAPI and mounted with Vectashield (Vector Labs). Image acquisition was performed on a Zeiss LSM800 AiryScan microscope (63X). A Python-based image processing workflow was developed to quantify colocalization across multiple samples. Each labeled nucleus was analyzed with region props to extract pixel intensities for downstream colocalization calculations. PearsonR and Manders’ coefficients (M1, M2) were computed for PCNA vs. GFP intensities. Analysis code is available at: https://zenodo.org/records/15465659.

### Proximity ligation assay (PLA)

Cells were fixed as described as for immunofluorescence. PLA processing was completed using the Navinci Diagnostics NaveniFlex MR100 PLA Kit (NF.MR.100.1) with primary antibodies (GFP 1:500, PCNA 1:500). Image acquisition was performed using a Zeiss Axio Imager Z1 (100X). Maximum intensity projections were generated for analysis (Image J).

### Capillary electrophoresis

Capillary electrophoresis of whole cell lysates was performed as described in Wittig et al.^28^. Samples were run on a Jess and analyzed using Compass Software (ProteinSimple). The following antibodies were used: CDT2 (Abcam ab184548); Cyclin A2 (BioLegend 644001); CDC45L (Proteintech 15678-1-AP).

### Statistics

Graphs and associated statistical tests were performed using GraphPad Prism.

## Data availability

All flow cytometry data is available on FigShare (https://doi.org/10.6084/m9.figshare.29242556).

## Acknowledgements

We thank the Bioinformatics and Pathways Core, supported by COBRE grant 5P30GM149376-02, for assistance with data analysis. Cell sorting was performed by the OMRF Flow Cytometry Core and imaging was performed by OMRF Imaging Core Facility. We are grateful to Hideo Nishitani for the pCMV-HA-Cdt2(WT)-3×FLAG^50^ plasmid and Alan Bradley for the pCMV-hyPBase^51^ plasmid. The following plasmids were obtained from Addgene: pMK289^52^ (mAID-mClover-NeoR, #72827; Masato Kanemaki), pBabe Puro osTIR1-9Myc^53^ (#80074; Andrew Holland), XLone-GFP^54^ (#96930; Xiaojun Lian), pX330_human CDK1^39^ (#118597; William Earnshaw), CDK1as_T2A_Zeo^39^ (#118596; William Earnshaw), and pCMV(CAT)T7-SB100^55^ (#34879; Zsuzsanna Izsvak).

## Author contributions

Conceptualization: MSH, CLS.

Methodology: MSH, CGS, KAW, TDN, KAB.

Investigation: MSH, CLS.

Formal analysis: MSH.

Writing – original draft: MSH, CLS.

Writing – review & editing: CGS, CLS.

Supervision: CLS.

## Funding

Funding provided by National Institutes of Health [R01GM121703] and The Presbyterian Health Foundation. MSH and TDN received support from the John and Mildred Carson PhD Scholarship Fund awarded for the OMRF Pre-doctoral Scholarship.

## Notes

### Competing Interest Statement

The authors have declared no competing interest.

