## Supplemental information for "Dynamic regulation of origin firing factors links CDK activity to dormant origin activation"

**CONTENTS:**

Supplementary figures: 2
- Supplementary Fig. 1. Time-dependent effects of WEE1 inhibition on chromatin-bound TRESLIN, MTBP, and MCM7 levels. 2
- Supplementary Fig. 2. Characterization of CDK-dependent regulation of TRESLIN and MTBP recruitment and CDK1as cell line validation. 3
- Supplementary Fig. 3. siRNA knockdown efficacy was demonstrated by flow cytometry, and the extent of knockdown was not affected by WEE1 inhibition. 4
- Supplementary Fig. 4. CDT2 overexpression prevents WEE1i-induced stabilization of CDT1 but not TRESLIN. 5
- Supplementary Fig. 5. The TRESLIN-8A mutant is degraded normally during S phase and responds to CDK modulation. 6
- Supplementary Fig. 6. Construction of CDC45-mClover knock-in line. 7
- Supplementary Fig. 7. TRESLIN and MTBP are required for WEE1i-induced increases in DNA synthesis in RPE-1 cells. 8

Supplementary tables: 9
- Supplementary table S1 (Chemicals): 9
- Supplementary Table S2 (Antibodies): 9
- Supplementary Table S3 (siRNAs): 9
- Supplementary Table S4 (Cell lines): 9
- Supplementary Table S4 (Plasmids): 11

Supplementary methods: 12
- Generation of stable cell lines: 12
- Plasmid sequences: 13

Supplementary references: 21

**SUPPLEMENTARY FIGURES:**

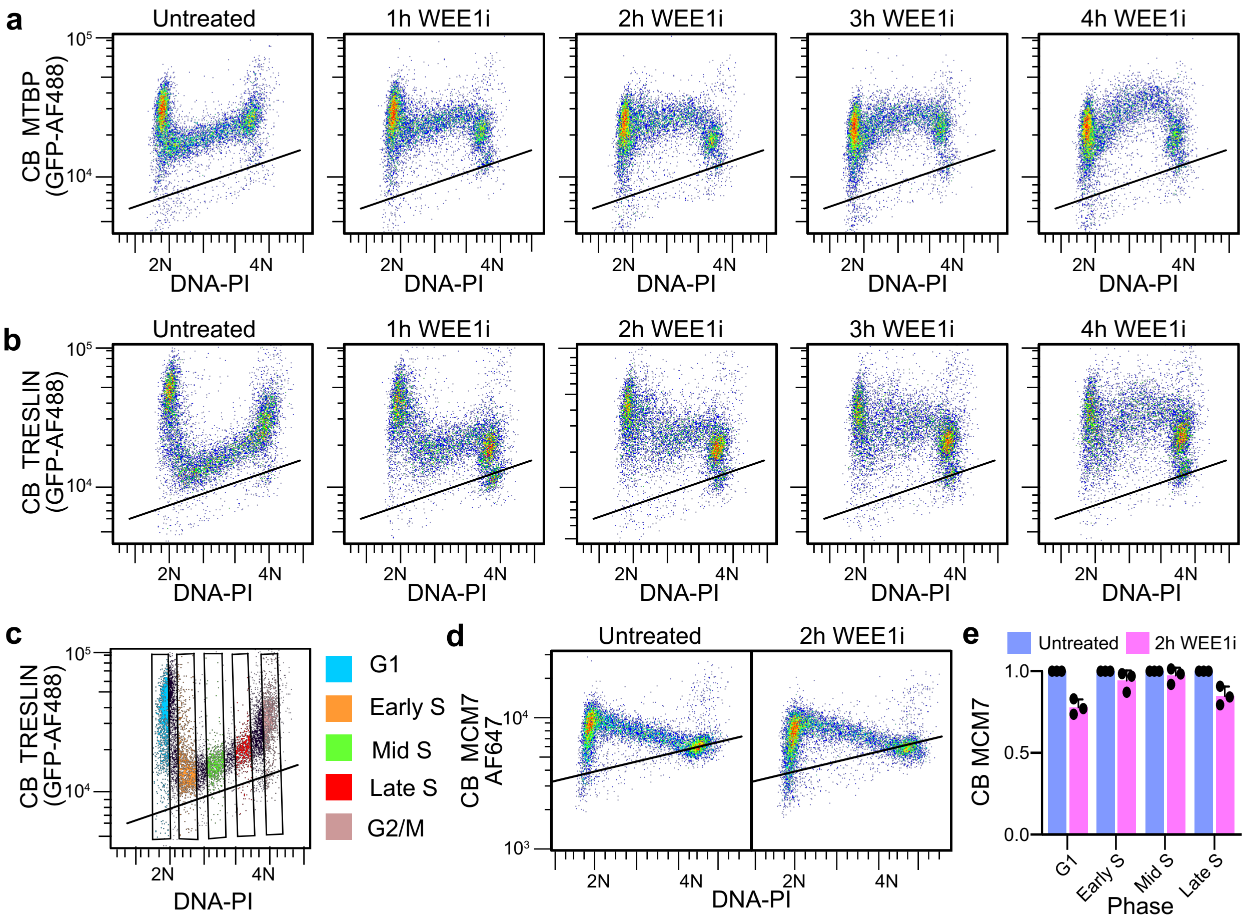


**Supplementary Fig. 1. Time-dependent effects of WEE1 inhibition on chromatin-bound TRESLIN, MTBP, and MCM7 levels. (a, b)** Flow cytometry analysis of chromatin-bound (CB) TRESLIN (a) and CB MTBP (b) in HCT116 cells in which endogenous TRESLIN or MTBP was tagged with mClover. Cells were extracted with CSK buffer to remove soluble proteins, immunolabeled with an anti-GFP antibody to detect the mClover tag, and stained with propidium iodide (PI) to measure DNA content. Pseudocolored dot plots show CB TRESLIN (a) or CB MTBP (b) (y-axis, log scale) as a function of DNA content (x-axis, linear scale). Five conditions are shown: untreated cells (leftmost plot) and cells treated with the WEE1 inhibitor (WEE1i; MK1775) for 1, 2, 3, or 4 hours. **(c)** Example of DNA content gates on dot plot of CB TRESLIN vs DNA content (propidium iodide) measured by flow cytometry. **(d)** Flow cytometry analysis of chromatin-bound MCM7 following WEE1i treatment. CB MCM7 levels were measured using an antibody against endogenous MCM7. Pseudocolored dot plots show CB MCM7 (y-axis, log scale) as a function of DNA content (x-axis, linear scale) in untreated cells (left) and cells treated with WEE1i for 2 hours (right). **(e)** The bar plot quantifies median CB MCM7 levels across the cell cycle, normalized to the median CB MCM7 in untreated G1 cells. Cells were binned into four DNA content groups: G1 (2N), early S (>2N), mid S (2N-4N), and late S (<4N). Bars represent mean values + SD of median values from three biological replicates.

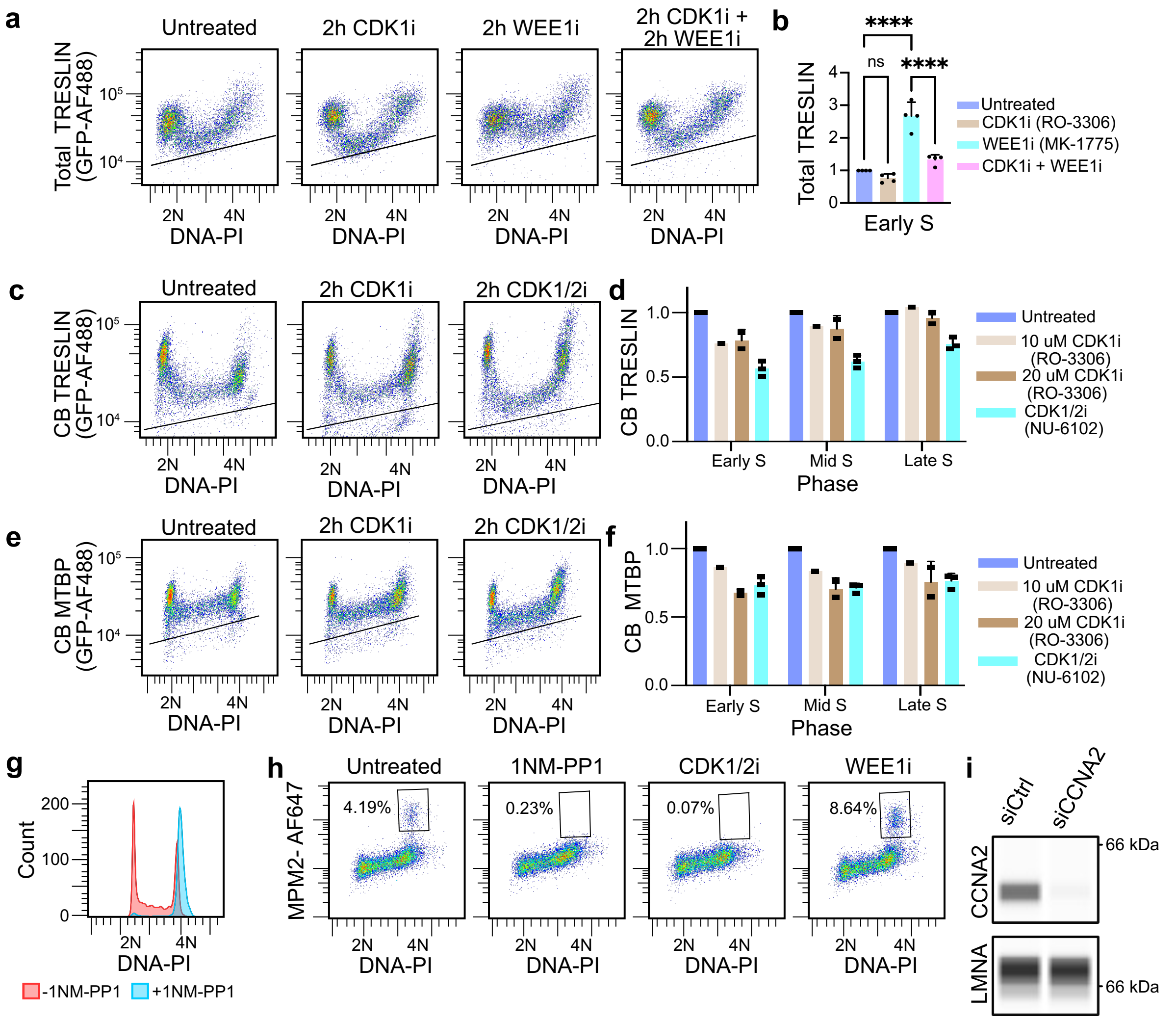


**Supplementary Fig. 2. Characterization of CDK-dependent regulation of TRESLIN and MTBP recruitment and CDK1as cell line validation.** **(a)** Representative pseudocolor dot plots showing flow cytometry measurement of total TRESLIN levels (anti-GFP signal) versus DNA content (propidium iodide; PI) in cells expressing endogenously mClover-tagged TRESLIN, treated with the indicated inhibitors. Black lines indicate background signal from untagged control cells stained in parallel. **(b)** Quantification of total TRESLIN levels from four biological replicates of the data shown in a. Median fluorescence was calculated for each cell cycle phase and sample, background-subtracted using the signal from untagged controls, and normalized to the untreated condition. **(c)** Same as a, but showing chromatin-bound (CB) TRESLIN instead of total TRESLIN, with the indicated inhibitor treatments. **(d)** Quantification of median CB-TRESLIN from three biological replicates of c, separated by early, mid, and late S-phase subpopulations. Values were background-subtracted and normalized to untreated within each S-phase fraction. **(e)** Same as c, but using cells expressing mClover-tagged MTBP to assess CB-MTBP levels. **(f)** Quantification of median CB-MTBP from three biological replicates of e, processed as described for d. **(g)** DNA content frequency (density) plot from flow cytometry of $CDK1^{as}$ cells treated with or without the ATP analog inhibitor 1NMPP1. **(h)** Pseudocolor dot plot showing flow cytometry measurement of anti-MPM2 (mitotic marker) versus DNA content in $CDK1^{as}$ cells treated with 1NMPP1. The boxed region indicates the 4N MPM2-positive mitotic population. The percentage of cells in the mitotic gate is shown on the plot. **(i)** Capillary electrophoresis (Jess) of Cyclin A2 (CCNA2) and LMNA loading control in whole-cell lysates from cells transfected with control siRNA or Cyclin A2-targeting siRNA, validating Cyclin A2 knockdown. In b statistical analysis was performed using one-way ANOVA followed by Tukey's multiple comparisons test. Significance levels: *p<0.05, **p<0.01, ***p<0.001, ****p<0.0001. Bar represents mean + SD of replicate medians.

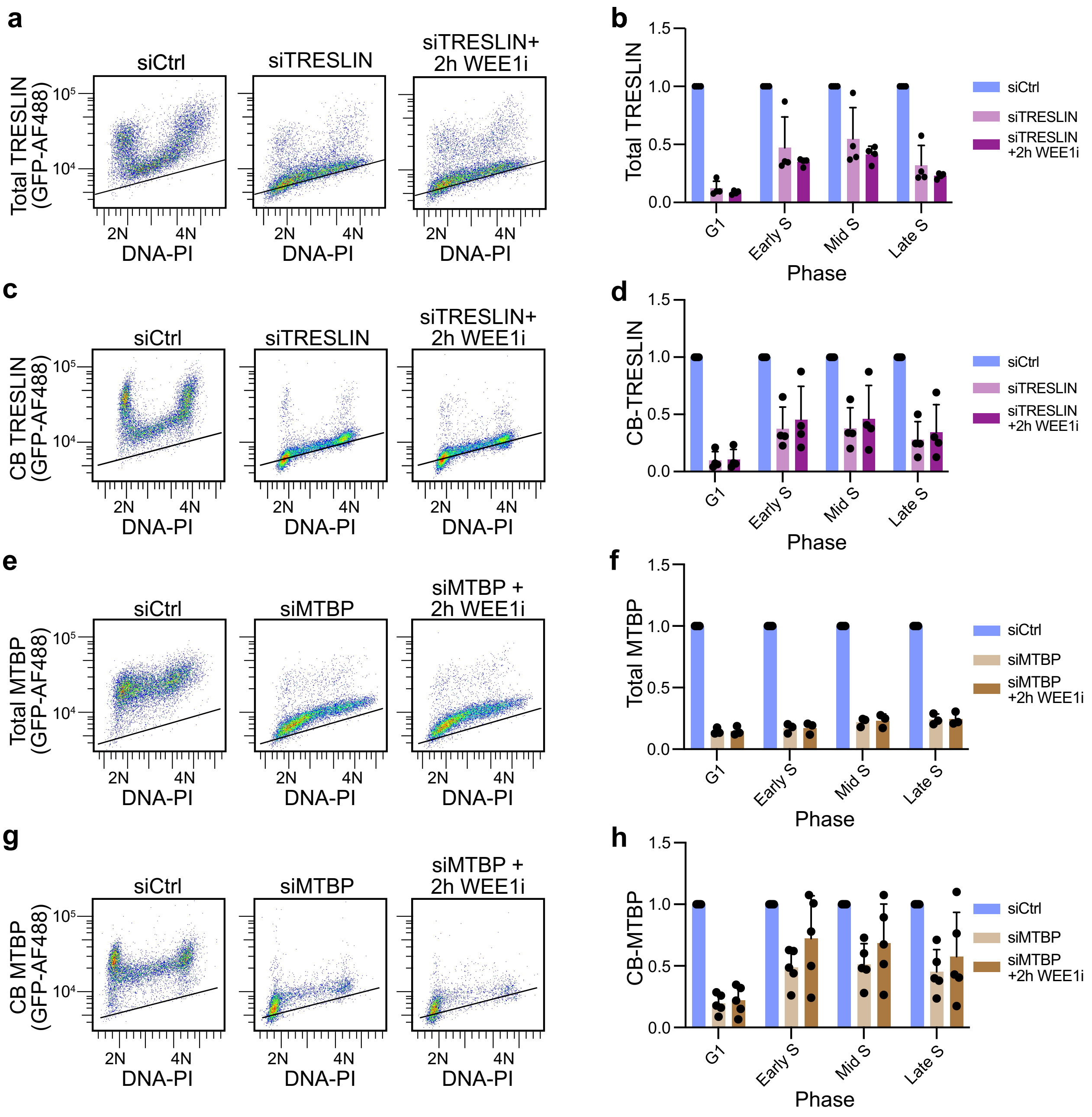


**Supplementary Fig. 3. siRNA knockdown efficacy was demonstrated by flow cytometry, and the extent of knockdown was not affected by WEE1 inhibition. (a, c, e, g)** Representative flow cytometry pseudocolor dot plots showing total (a, e) or chromatin-bound (CB) (c, g) levels of TRESLIN (a, c) or MTBP (e, g) versus DNA content (PI) in HCT116 cells expressing mClover-tagged endogenous TRESLIN or MTBP. Cells were transfected with non-targeting control siRNA (siCtrl), TRESLIN-targeting siRNA (siTRESLIN), or MTBP-targeting siRNA (siMTBP), and treated ± WEE1 inhibitor (WEE1i). A black line overlays the background GFP signal measured in parallel from untagged parental cells processed identically. **(b, d, f, h)** Quantification of flow cytometry data shown in a, c, e, and g, respectively. Bar plots represent background-subtracted and siCtrl-normalized median anti-GFP signal from individual replicates ($n \geq 3$), stratified by cell cycle stage (G1, early S, mid S, late S) based on DNA content. Data are presented as mean + SD of replicate medians.

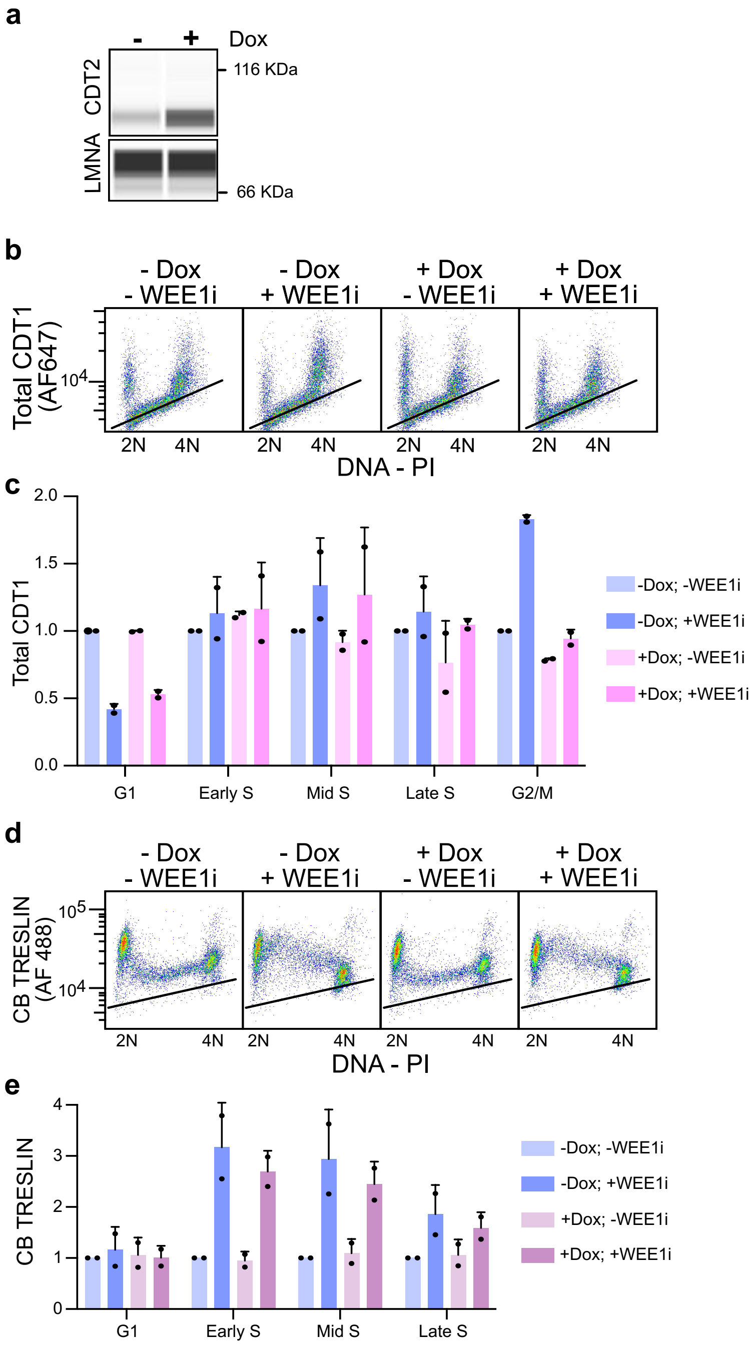


**Supplementary Fig. 4. CDT2 overexpression prevents WEE1i-induced stabilization of CDT1 but not TRESLIN. (a)** Capillary electrophoresis (Jess) of whole-cell lysates from HCT116 cells with a stable piggyBac Tet-On transgene encoding CDT2. Cells were treated with or without doxycycline (Dox) to induce CDT2 expression. CDT2 was detected with an anti-CDT2 antibody, and LMNA was used as a loading control. **(b)** Flow cytometry analysis of CDT1 levels in cells treated with Dox and/or WEE1 inhibitor (WEE1i; MK1775). CDT1 (y-axis, log scale) was detected using an anti-CDT1 antibody; DNA content (x-axis, linear scale) was measured by propidium iodide (PI) staining. The black line shows background signal from a control lacking the primary antibody. **(c)** Quantification of background-subtracted CDT1 signal from two biological replicates. Median values were calculated for cells in G1, early S, mid S, late S, and G2/M, and normalized within each stage to the –Dox/–WEE1i control. Bars show mean ± range. **(d)** Flow cytometry analysis of chromatin-bound (CB) TRESLIN in HCT116 cells with endogenous TRESLIN tagged with mClover, after Dox and/or WEE1i treatment. Black line represents background from untagged cells processed in parallel. **(e)** Quantification of background-subtracted CB TRESLIN signal from two biological replicates, calculated as in (c).

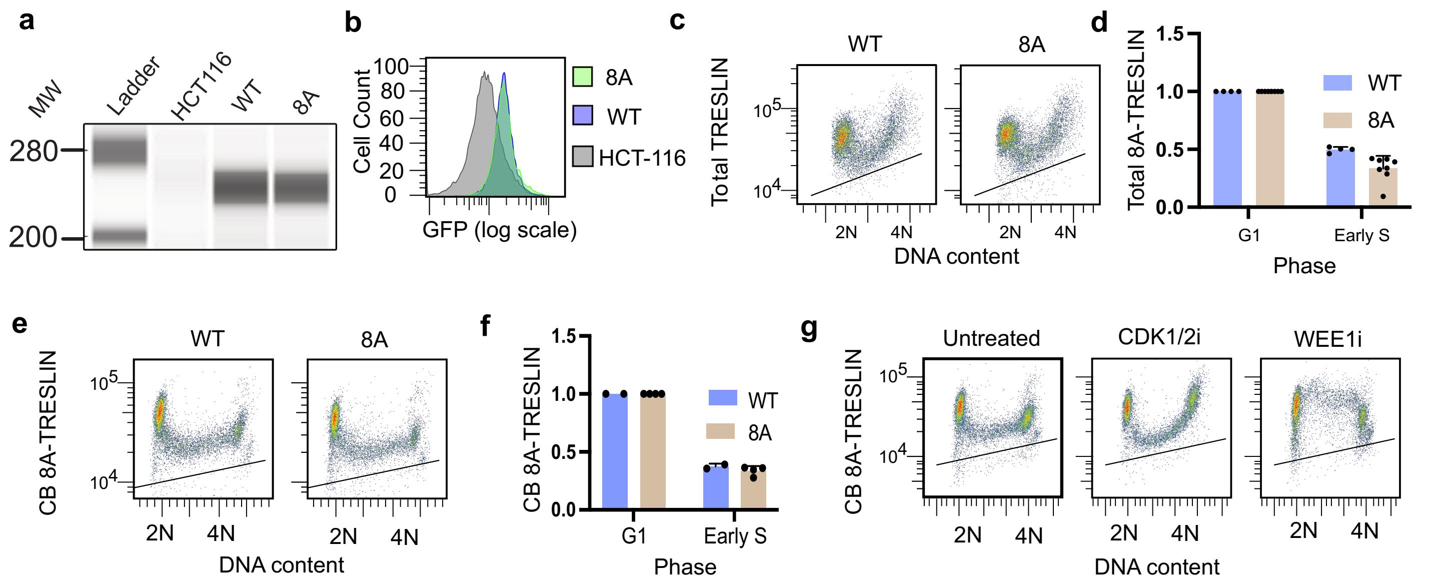


**Supplementary Fig. 5. The TRESLIN-8A mutant is degraded normally during S phase and responds to CDK modulation.** **(a)** Capillary electrophoresis (Jess) of whole-cell lysates from HCT116 cells with CRISPR knock-in of the TRESLIN-8A mutant, in which eight conserved charged residues in the SBI region were mutated. GFP-tagged TRESLIN was detected using an anti-GFP antibody. **(b)** Live-cell flow cytometry showing GFP fluorescence in TRESLIN-8A knock-in cells. Frequency histograms display cell counts versus GFP intensity (log scale). **(c)** Flow cytometry of total TRESLIN levels in TRESLIN-8A-mClover cells. Anti-GFP signal (log scale) is plotted against DNA content (PI, linear scale). **(d)** Quantification of total TRESLIN from (c) across replicates. Median anti-GFP signal was background-subtracted and normalized to the G1 phase signal within each replicate. **(e)** Flow cytometry of chromatin-bound (CB) TRESLIN-8A, plotted as in (c). **(f)** Quantification of CB TRESLIN from (e), performed as in (d). **(g)** Flow cytometry analysis of CB TRESLIN-8A levels in cells treated for 2 hours with CDK1/2 inhibitor (CDK1/2i; NU6102) or WEE1 inhibitor (WEE1i; MK1775), as in (e).

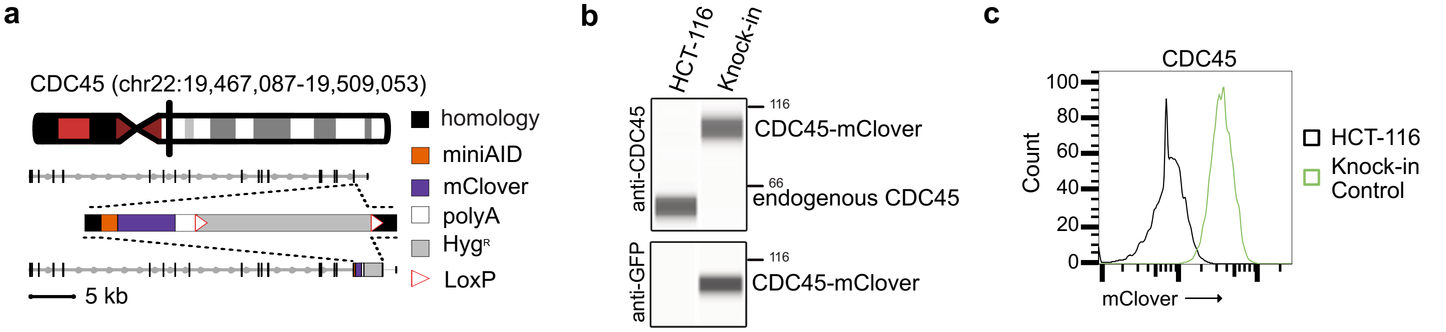


**Supplementary Fig. 6. Construction of CDC45-mClover knock-in line. (a)** Schematic of the chromosome ideogram, targeting location, and targeting constructs for C-terminal tagging of CDC45 with a miniAID-mClover tag. Homology arms are ~200bp. **(b)** Capillary electrophoresis (Jess) of whole-cell lysates from HCT116 cells with CRISPR knock-in of mini-AID-mClover tag into the *CDC45* gene. Total CDC45 was detected with anti-CDC45 antibody. Size shift is consistent with the addition of tag to all alleles. GFP-tagged CDC45 was detected using an anti-GFP antibody. **(c)** Live-cell flow cytometry showing GFP fluorescence in CDC45 knock-in cells. Frequency histograms display cell counts versus GFP intensity (log scale).

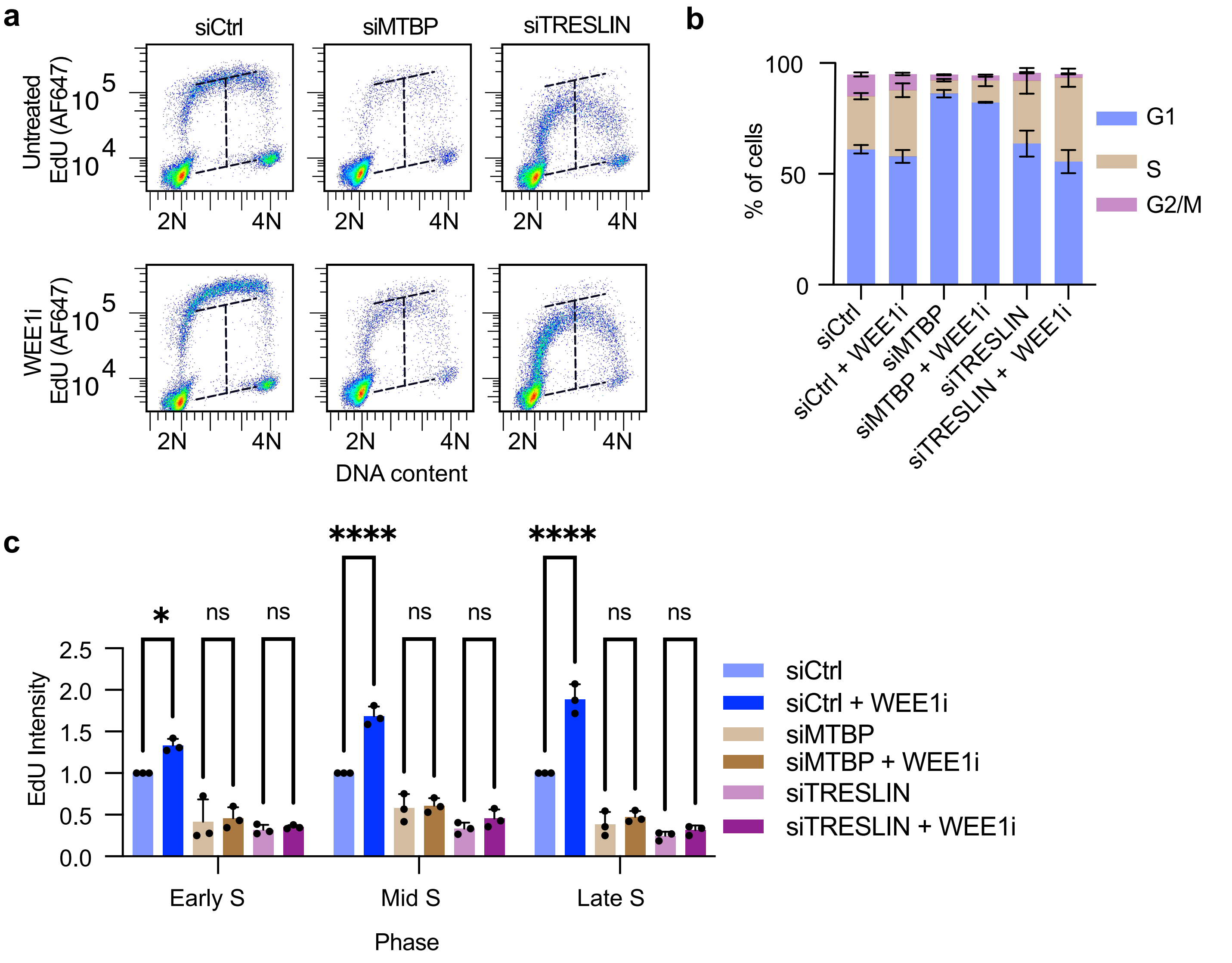


**Supplementary Fig. 7. TRESLIN and MTBP are required for WEE1i-induced increases in DNA synthesis in RPE-1 cells. (a)** Representative flow cytometry pseudocolor plots showing EdU incorporation versus DNA content (propidium iodide) in RPE-1 cells transfected with siControl, siTRESLIN, or siMTBP, with or without WEE1 inhibitor (WEE1i) treatment. **(b)** Stacked bar plots showing the proportion of cells in G1, S, or G2/M phases for each siRNA and treatment condition, based on EdU versus DNA flow cytometry data from a (n = 3 biological replicates). Bars represent means and error bars represent standard deviation (SD). **(c)** Quantification of EdU intensity in early, mid, and late S phase fractions from the experiment in a, based on DNA content gating. Values represent median EdU signal per cell, background subtracted and normalized to siControl within each S phase fraction. Bars show means and error bars represent SD from three biological replicates. Statistical comparisons were performed using two-way ANOVA with Tukey’s post-hoc test.

**SUPPLEMENTARY TABLES:**

**Supplementary table S1 (Chemicals):**

| Chemical | Concentration | Catalog Number | Source |
|---|---|---|---|
| MLN4924 | 3uM | 15217 | Cayman Chemical |
| NU-6102 | 20 uM | 13317 | Cayman Chemical |
| Doxycycline | 2.5 ug/ml | 380-273-g005 | Enzo |
| MK-1775 | 1uM | 21266 | Selleck Chemicals |
| RO3306 | 10 uM | 15149 | Cayman Chemical |
| PF-06873600 | 25 nM | 35502 | Cayman Chemical |
| 1 NM-PP1 | 5uM | 529581 | Calbiochem |
| EdU | 20uM | E1018 | Life Technologies |

**Supplementary Table S2 (Antibodies):**

| Antibody | Catalog Number | Lot Number | Source |
|---|---|---|---|
| GFP | 600-401-215S | 43570 | Rockland |
| PCNA | Sc-56 | E2418 | Santa Cruz |
| TRESLIN | A303-472A | 1 | Bethyl |
| CDT1 | ab202067 | GR329107-2 | Abcam |
| MPM2 | M3514 | 124 | DAKO |
| MCM7 | Sc-56324 | B0718 | Santa Cruz |
| Cyclin A2 | 644001 | B273965 | BioLegend |
| CDC45L | 15678-1-AP | 00011415 | ProteinTech |
| CDT2 | Ab18458 | GR3233757-2 | Abcam |
| Lamin A/C | 2032S | 6 | Cell Signaling |

**Supplementary Table S3 (siRNAs):**

| siRNA | ID | Source |
|---|---|---|
| siCyclin A2 | S2512 | Life Technologies |
| siTreslin | Kumagai et al. 2010[1] | Life Technologies |
| siMTBP | M-013953–01-0010 | Dharmacon |
| siCtrl | 12935110 | Life Technologies |
| siPCNA | VC30004 | Sigma Aldrich |

**Supplementary Table S4 (Cell lines):**

| Cell line | Parental Cell Line | Genotype | Source | RRID |
|---|---|---|---|---|
| HCT116 | | | ATCC (CCL-247) | RRID:CVCL_0291 |
| HCT-116 MTBP-mClover | HCT116 | | Sansam lab; Wittig et al[2] | RRID:CVCL_C7SR |
| HCT-116 TICRR-mClover | HCT116 | | Sansam lab; Wittig et al[2] | RRID:CVCL_C7SS |

| | | | | |
|---|---|---|---|---|
| Flp-In-T-REx-293 | | | Thermo Fisher (R78007) | RRID:CVCL_U427 |
| HCT-116 CDC45-mClover | HCT116 | biallelic knock-in of a miniAID–mClover–polyA cassette followed by a PGK-HygroR transgene, inserted just upstream of the CDC45 stop codon | Sansam lab; This study; donor plasmid: pBSKII+_hCDC45 mClover-mAID HygroR; CRISPR: pX330- | Pending |
| HCT116 TICRR-mClover;TRE3G::CDT2;EF1a::rtTA3 | HCT116 TICRR-mClover | HCT116 cell line with an mClover knock-in at the endogenous TICRR locus, and a doxycycline-inducible CDT2 transgene under the TRE3G promoter. rtTA3 is constitutively expressed from the EF1α promoter to support Tet-On regulation. | Sansam lab; This study | Pending |
| HCT116 TICRR-GT[SA–mClover– ex20–22(WT)] | HCT116 | HCT116 cell line with a gene trap targeted to intron 19 of TICRR, containing a strong splice acceptor, followed by an in-frame mClover-tagged TICRR coding region (exons 20–22). The cassette includes a polyadenylation signal and a PGK promoter-driven Hygromycin resistance gene for selection. | Sansam lab; This study | Pending |
| HCT116 TICRR-GT[SA–mClover– ex20–22(8A)] ex20–22(WT/ΔSBI/8A) | HCT116 | HCT116 cell line with a gene trap targeted to intron 19 of TICRR, containing a strong splice acceptor, followed by an in-frame mClover-tagged TICRR coding region (exons 20–22) with 8A mutations. The cassette includes a polyadenylation signal and a PGK promoter-driven Hygromycin resistance gene for selection. | Sansam lab; This study | Pending |
| HCT116 TICRR-GT[SA–mClover– ex20–22(ΔSBI)] | HCT116 | HCT116 cell line with a gene trap targeted to intron 19 of TICRR, containing a strong splice acceptor, followed by an in-frame mClover-tagged TICRR coding region (exons 20–22) with ΔSBI deletion. The cassette includes a polyadenylation signal and a PGK promoter-driven Hygromycin resistance gene for selection. | Sansam lab; This study | Pending |

**Supplementary Table S4 (Plasmids):**

| Plasmid | Source | Notes |
|---|---|---|
| pBSKII+_hCDC45 mClover-mAID HygroR | Sansam lab; This study | Targeting vector for C-terminal CDC45 knockin; See sequence below |
| pBlueScriptIISKPlus_TICRRIntronTargeting3_TICRR_WT (9B8) | Sansam lab; This study | Targeting vector for intronic TICRR gene trap knockin; See sequence below |
| pBlueScriptIISKPlus_TICRRIntronTargeting3_TICRR_8A (9C1) | Sansam lab; This study | Targeting vector for intronic TICRR gene trap knockin; See sequence below |
| pBlueScriptIISKPlus_TICRRIntronTargeting3_TICRR_ΔSBI (9B9) | Sansam lab; This study | Targeting vector for intronic TICRR gene trap knockin; See sequence below |
| pX330-TICRR(CS2391) | Sansam lab; This study | Oligos: 5'-CAC CGA ATG TGA TTG GTG CAG TGA C-3'; 5'-AAA CGT CAC TGC ACC AAT CAC ATT C-3' |
| pX330-TICRR(CS2389) | Sansam lab; This study | Oligos: 5'-CAC CGC AGT GAC AGG GAC ATG CGA G-3'; 5'-AAA CCT CGC ATG TCC CTG TCA CTG C-3' |
| pX330-hCDC45(CS1997) | Sansam lab; This study | Oligos: 5'-CAC CGG TCC TAG GGT GAG TTA CAG-3'; 5'-AAA CCT GTA ACT CAC CCT AGG ACC-3' |
| pT2/SVNeo | Addgene 26553 – Cui et al., 2002[3] | |
| pCMV(CAT)T7-SB100 | Addgene 34879 – Mates et al., 2009[4] | |
| pX330-U6_Chimeric_BB-cBh-hSpCas9 | Addgene 42230 – Cong et al, 2013[5] | |
| pX330_human CDK1 | Addgene 118597 – Saldivar et al, 2018[6] | |
| CDK1as_T2A_Zeo | Addgene 118596 - Saldivar et al, 2018[6] | |
| pCYL43 | Wellcome Trust Sanger Institute | PiggyBac transposase |
| XLone-GFP | Addgene 96930 – | |

| | | |
|---|---|---|
| | Randolph, et al, 2017[7] | |
| XLone-CDT2 | Sansam lab; This study | PiggyBac Transposon for Tet-inducible CDT2; See sequence below |

**SUPPLEMENTARY METHODS:**

**Generation of stable cell lines:**

**HCT-116 CDC45-mClover**

The HCT116 CDC45-mClover knock-in cell line was generated using pX330 sgRNA expression (pX330-hCDC45(CS1997)) and donor (pBSKII+_hCDC45 mClover-mAID HygroR) plasmids for CDC45 following the procedure described by Wittig et al.[2].

**HCT-116 MTBP-mClover;CDK1-/-;CMV::xCDK1$^{AS}$**
**HCT-116 TICRR-mClover;CDK1-/-;CMV::xCDK1$^{AS}$**

For the creation of CDK1 analog sensitive lines, the one-shot method described by Saldivar et al was used. Briefly, HCT-116 MTBP-mClover and HCT-116 TICRR-mClover were cotransfected with pCMV(CAT)T7-SB100, CDK1as_T2A_Zeo, and pX330_human CDK1 and selected with zeocin before dilution cloning. Surviving clones were tested with 1NM-PP1 treatment to identify those with inhibited mitotic entry.

**HCT116 TICRR-mClover;TRE3G::CDT2;EF1a::rtTA3**

To generate the doxycycline-inducible CDT2 transgenic line, HCT116 TICRR-mClover cells were co-transfected with two plasmids: XLone-CDT2, containing CDT2 under the control of the TRE3G promoter and the Tet-On 3G transactivator driven by the EF1α promoter, and pCYL43, encoding a hyperactive PiggyBac transposase. Following transfection, cells were selected with blasticidin, and clonal lines were screened for doxycycline-inducible CDT2 expression by capillary electrophoresis.

**HCT116 TICRR-GT[SA–mClover– ex20–22(WT)]**
**HCT116 TICRR-GT[SA–mClover– ex20–22(ΔSBI)]**
**HCT116 TICRR-GT[SA–mClover– ex20–22(8A)]**

To generate the intronic gene trap lines HCT116-WT-TRESLIN, HCT116-ΔSBI-TRESLIN, and HCT116-8A-TRESLIN, a homology-directed repair (HDR) strategy was used to insert donor exons into an intronic region between exons 19 and 20 of the endogenous TICRR locus. Each donor construct included a splice acceptor followed by exons 20–22, an mClover fusion at the C-terminus, a stop codon, and a polyadenylation signal. Three variants were used: WT-TRESLIN (wild-type sequence), ΔSBI-TRESLIN (deletion of residues 1485–1661), and 8A-TRESLIN (mutation of residues 1561–1580 from EVELEMQASGLPKLRIKKID to AVALAMQASGLPALAIAAIA). CRISPR/Cas9 was used to introduce double-strand breaks at the intronic target site. Clones were selected with 200 µg/mL hygromycin, screened by PCR, and validated for expression of the tagged protein by capillary electrophoresis using anti-GFP and anti-TRESLIN antibodies.

**293 Flp-In T-REx-CMV/TetO2::mfGFP-TRESLIN**
**293 Flp-In T-REx-CMV/TetO2::mfGFP-TRESLIN(1-1111)**
**293 Flp-In T-REx-CMV/TetO2::mfGFP-TRESLIN(1059-1910)**

293 Flp-In T-REx cell lines expressing N-terminal or C-terminal fragments of TRESLIN were generated by co-transfecting cells with the Flp recombinase plasmid pOG44 (Invitrogen) and mfGFP-TRESLIN expression constructs in pcDNA5/FRT/TO using TransIT-LT1 (Mirus Bio). The N-terminal (residues 1–1111) and C-terminal (residues 1059–1910) fragments of TRESLIN were cloned into the pcDNA5/FRT/TO vector (Invitrogen) via isothermal assembly. Stable clones were selected using 200 µg/mL hygromycin.

**Plasmid sequences:**

>pBSKII+_hCDC45 mClover-mAID HygroR (7C3)
atccggtgcaggcgccaaggagaagagtgcttgtcctaaagatccagccaaacctccggccaaggcacaagttgtgggatggccaccggtgagatcat
accggaagaacgtgatggtttcctgccaaaaatcaagcggtggcccggaggcggcggcgttcgtgaaggtatcaatggacggagcaccgtacttgagg
aaaatcgatttgaggatgtataaagctagcatggtgagcaagggcgaggagctgttcaccggggtggtgcccatcctggtcgagctggacggcgacgtaa
acggccacaagttcagcgtccgcggcgagggcgagggcgatgccaccaacggcaagctgaccctgaagttcatctgcaccaccggcaagctgcccgt
gccctggcccaccctcgtgaccaccttcggctacggcgtggcctgcttcagccgctaccccgaccacatgaagcagcacgacttcttcaagtccgccatgc
ccgaaggctacgtccaggagcgcaccatctctttcaaggacgacggtacctacaagacccgcgccgaggtgaagttcgagggcgacaccctggtgaac
cgcatcgagctgaagggcatcgacttcaaggaggacggcaacatcctggggcacaagctggagtacaacttcaacagccacaacgtctatatcacggc
cgacaagcagaagaacggcatcaaggctaacttcaagatccgccacaacgttgaggacggcagcgtgcagctcgccgaccactaccagcagaacac
ccccatcggcgacggccccgtgctgctgcccgacaaccactacctgagccatcagtccaagctgagcaaagaccccaacgagaagcgcgatcacatg
gtcctgctggagttcgtgaccgccgccgggattacacatggcatggacgagctgtacaagtaatctagataactgatcataatcagccataccacatttgtag
aggttttacttgctttaaaaaacctcccacacctccccctgaacctgaaacataaaatgaatgcaattgttgttgttaacttgtttattgcagcttataatggttacaa
ataaagcaatagcatcacaaatttcacaaataaagcatttttttcactgcattctagttgtggtttgtccaaactcatcaatgtatcttaacgcgtcgatcatattcaa
taacccttaatataacttcgtataatgtatgctatacgaagttattaggtctgaagaggagtttacgtccagccaagcttaggatctcgacctcgaaattctaccg
ggtaggggaggcgcttttcccaaggcagtctggagcatgcgctttagcagccccgctgggcacttggcgctacacaagtggcctctggcctcgcacacattc
cacatccaccggtaggcgccaaccgactccgttctttggtggccccttcgcgccaccttctactcctcccctagtcaggaagttcccccccgccccgcagctc
gcgtcgtgcaggacgtgacaaatggaagtagcacgtctcactagtctcgtgcagatggacagcaccgctgagcaatggaagcgggtaggcctttggggc
agcggccaatagcagctttgctccttcgctttctgggctcagaggctgggaaggggtgggtccgggggcgggctcaggggcgggctcaggggcggggcg
ggcgcccgaaggtcctccggaggcccggcattctgcacgcttcaaaagcgcacgtctgccgcgctgttctcctcttcctcatctccgggcctttcgacctgcat
ccatctagatctcgagcagctgaagcttaccgctagcatggatagatccggaaagcctgaactcaccgcgacgtctgtcgagaagtttctgatcgaaaagtt
cgacagcgtctccgacctgatgcagctctcggagggcgaagaatctcgtgctttcagcttcgatgtaggagggcgtggatatgtcctgcgggtaaatagctg
cgccgatggtttctacaaagatcgttatgtttatcggcactttgcatcggccgcgctcccgattccggaagtgcttgacattggggaattcagcgagagcctgac
ctattgcatctcccgccgtgcacagggtgtcacgttgcaagacctgcctgaaaccgaactgcccgctgttctgcagccggtcgcggaggccatggatgcgat
cgctgcggccgatcttagccagacgagcgggttcggcccattcggaccgcaaggaatcggtcaatacactacatggcgtgatttcatatgcgcgattgctga
tccccatgtgtatcactggcaaactgtgatggacgacaccgtcagtgcgtccgtcgcgcaggctctcgatgagctgatgctttgggccgaggactgccccga
agtccggcacctcgtgcacgcggatttcggctccaacaatgtcctgacggacaatggccgcataacagcggtcattgactggagcgaggcgatgttcggg
gattcccaatacgaggtcgccaacatcttcttctggaggccgtggttggcttgtatggagcagcagacgcgctacttcgagcggaggcatccggagcttgca
ggatcgccgcggctccgggcgtatatgctccgcattggtcttgaccaactctatcagagcttggttgacggcaatttcgatgatgcagcttgggcgcagggtcg
atgcgacgcaatcgtccgatccggagccgggactgtcgggcgtacacaaatcgcccgcagaagcgcggccgtctggaccgatggctgtgtagaagtact
cgccgatagtggaaaccgacgccccagcactcgtccgagggcaaaggaataggctagccgcccgccccacgacccgcagcgcccgaccgaaagga
gcgcacgaccccatgcatcgatgatatcagatccccgggatgcagaaattgatgatctattaaacaataaagatgtccactaaaatggaagtttttcctgtcat
actttgttaagaagggtgagaacagagtacctacattttgaatggaaggattggagctacgggggtgggggtggggtgggattagataaatgcctgctcttta
ctgaaggctctttactattgctttatgataatgtttcatagttggatatcataatttaaacaagcaaaaccaaattaagggccagctcattcctcccactcatgatct
atagatctatagatctctcgtgggatcattgtttttctcttgattcccactttgtggttctaagtactgtggtttccaaatgtgtcagtttcatagcctgaagaacgagat
cagcagcctctgttccacatacacttcattctcagtattgttttgccaagttctaattccatcagaagctggtcgagatccggaacccttaatataacttcgtataat
gtatgctatacgaagttattaggtccctcgaagaggttcactagtagggtgagttacaggggttctgcaggggtggctgcagcagcccccctcagagcccgac
cctgatgccctgctctgtcctccctcaacggaggcttctacttgggtttcagaccgaagcagggttcttgagattggagccaacacatttttcccaagcacatctg
tccttaggctgccagcagggccacaatggggcatttatcaagcttatcgataccgtcgacctcgagggggggcccggtacccaattcgccctatagtgagtc
gtattacgcgcgctcactggccgtcgttttacaacgtcgtgactgggaaaaccctggcgttacccaacttaatcgccttgcagcacatccccctttcgccagct
ggcgtaatagcgaagaggcccgcaccgatcgcccttcccaacagttgcgcagcctgaatggcgaatggaaattgtaagcgttaatattttgttaaaattcgc
gttaaatttttgttaaatcagctcattttttaaccaataggccgaaatcggcaaaatcccttataaatcaaaagaatagaccgagatagggttgagtgttgttcca
gtttggaacaagagtccactattaaagaacgtggactccaacgtcaaagggcgaaaaaccgtctatcagggcgatggcccactacgtgaaccatcaccct
aatcaagttttttggggtcgaggtgccgtaaagcactaaatcggaaccctaaagggagcccccgatttagagcttgacggggaaagccggcgaacgtgg
cgagaaaggaagggaagaaagcgaaaggagcgggcgctagggcgctggcaagtgtagcggtcacgctgcgcgtaaccaccacacccgccgcgctt
aatgcgccgctacagggcgcgtcaggtggcacttttcggggaaatgtgcgcggaacccctatttgtttatttttctaaatacattcaaatatgtatccgctcatga
gacaataaccctgataaatgcttcaataatattgaaaaaggaagagtatgagtattcaacatttccgtgtcgcccttattcccttttttgcggcattttgccttcctgtt
tttgctcacccagaaacgctggtgaaagtaaaagatgctgaagatcagttgggtgcacgagtgggttacatcgaactggatctcaacagcggtaagatcctt
gagagttttcgccccgaagaacgttttccaatgatgagcacttttaaagttctgctatgtggcgcggtattatcccgtattgacgccgggcaagagcaactcggt
cgccgcatacactattctcagaatgacttggttgagtactcaccagtcacagaaaagcatcttacggatggcatgacagtaagagaattatgcagtgctgcc
ataaccatgagtgataacactgcggccaacttacttctgacaacgatcggaggaccgaaggagctaaccgcttttttgcacaacatgggggatcatgtaact
cgccttgatcgttgggaaccggagctgaatgaagccataccaaacgacgagcgtgacaccacgatgcctgtagcaatggcaacaacgttgcgcaaacta

ttaactggcgaactacttactctagcttcccggcaacaattaatagactggatggaggcggataaagttgcaggaccacttctgcgctcggcccttccggctg
gctggtttattgctgataaatctggagccggtgagcgtgggtctcgcggtatcattgcagcactggggccagatggtaagccctcccgtatcgtagttatctaca
cgacggggagtcaggcaactatggatgaacgaaatagacagatcgctgagataggtgcctcactgattaagcattggtaactgtcagaccaagtttactca
tatatactttagattgatttaaaacttcatttttaatttaaaaggatctaggtgaagatcctttttgataatctcatgaccaaaatcccttaacgtgagttttcgttccact
gagcgtcagaccccgtagaaaagatcaaaggatcttcttgagatcctttttttctgcgcgtaatctgctgcttgcaaacaaaaaaaccaccgctaccagcggt
ggtttgtttgccggatcaagagctaccaactctttttccgaaggtaactggcttcagcagagcgcagataccaaatactgttcttctagtgtagccgtagttaggc
caccacttcaagaactctgtagcaccgcctacatacctcgctctgctaatcctgttaccagtggctgctgccagtggcgataagtcgtgtcttaccgggttggac
tcaagacgatagttaccggataaggcgcagcggtcgggctgaacggggggttcgtgcacacagcccagcttggagcgaacgacctacaccgaactga
gatacctacagcgtgagctatgagaaagcgccacgcttcccgaagggagaaaggcggacaggtatccggtaagcggcagggtcggaacaggagag
cgcacgagggagcttccagggggaaacgcctggtatctttatagtcctgtcgggtttcgccacctctgacttgagcgtcgatttttgtgatgctcgtcaggggggc
ggagcctatggaaaaacgccagcaacgcggcctttttacggttcctggccttttgctggccttttgctcacatgttctttcctgcgttatcccctgattctgtggata
accgtattaccgcctttgagtgagctgataccgctcgccgcagccgaacgaccgagcgcagcgagtcagtgagcgaggaagcggaagagcgcccaat
acgcaaaccgcctctccccgcgcgttggccgattcattaatgcagctggcacgacaggtttcccgactggaaagcgggcagtgagcgcaacgcaattaat
gtgagttagctcactcattaggcaccccaggctttacactttatgcttccggctcgtatgttgtgtggaattgtgagcggataacaatttcacacaggaaacagct
atgaccatgattacgccaagcgcgcaattaaccctcactaaagggaacaaaagctggagctccaccgcggtggcggccgctctagaactagtggatccc
ccgggctgcaggaattcgatgagtgcagaataccacaggccgggaggagccgcgcactttggaatgcagtggagggcaggcagcggaggggagttct
gtgccctgtcttgtgttcccactcctcccttctcacggctgtttttctttcattacttcagtaattgagctgaaagctgaggatcggagcaagtttctggacgcacttatt
tccctcctgtccgg

>pBlueScriptIISKPlus_TICRRIntronTargeting3_TICRR_WT (9B8)
atgtcctgcgggtaaatagctgcgccgatggtttctacaaagatcgttatgtttatcggcactttgcatcggccgcgctcccgattccggaagtgcttgacattgg
ggaattcagcgagagcctgacctattgcatctcccgccgtgcacagggtgtcacgttgcaagacctgcctgaaaccgaactgcccgctgttctgcagccggt
cgcggaggccatggatgcgatcgctgcggccgatcttagccagacgagcgggttcggcccattcggaccgcaaggaatcggtcaatacactacatggcg
tgatttcatatgcgcgattgctgatccccatgtgtatcactggcaaactgtgatggacgacaccgtcagtgcgtccgtcgcgcaggctctcgatgagctgatgct
ttgggccgaggactgccccgaagtccggcacctcgtgcacgcggatttcggctccaacaatgtcctgacggacaatggccgcataacagcggtcattgact
ggagcgaggcgatgttcggggattcccaatacgaggtcgccaacatcttcttctggaggccgtggttggcttgtatggagcagcagacgcgctacttcgagc
ggaggcatccggagcttgcaggatcgccgcggctccgggcgtatatgctccgcattggtcttgaccaactctatcagagcttggttgacggcaatttcgatgat
gcagcttgggcgcagggtcgatgcgacgcaatcgtccgatccggagccgggactgtcgggcgtacacaaatcgcccgcagaagcgcggccgtctgga
ccgatggctgtgtagaagtactcgccgatagtggaaaccgacgccccagcactcgtccgagggcaaaggaataggctagccgcccgccccacgaccc
gcagcgcccgaccgaaaggagcgcacgaccccatgcatcgatgatatcagatccccgggatgcagaaattgatgatctattaaacaataaagatgtcca
ctaaaatggaagtttttcctgtcatactttgttaagaagggtgagaacagagtacctacattttgaatggaaggattggagctacgggggtgggggtggggtg
ggattagataaatgcctgctctttactgaaggctctttactattgctttatgataatgtttcatagttggatatcataatttaaacaagcaaaaccaaattaagggcc
agctcattcctcccactcatgatctatagatctatagatctctcgtgggatcattgtttttctcttgattcccactttgtggttctaagtactgtggtttccaaatgtgtcag
tttcatagcctgaagaacgagatcagcagcctctgttccacatacacttcattctcagtattgttttgccaagttctaattccatcagaagctggtcgagatccgga
acccttaatataacttcgtataatgtatgctatacgaagttattaggtccctcgaagaggttcactaggatccgcatgtccctgtcactgcaccaatcacattgtat
tttcacctgcgtatgccccactgaacaggaaatgccagctagactgagagtcccctgacgggatgaaggaaggctgtaggaaagataagcattgatttcctt
acctgttggagcttttcactacgtatttttgcaagaaggcagggaaatttttttattggggagtgtcttttctgaatcaagcttatcgataccgtcgacctcgagggggg
ggcccggtacccaattcgccctatagtgagtcgtattacgcgcgctcactggccgtcgttttacaacgtcgtgactgggaaaaccctggcgttacccaacttaa
tcgccttgcagcacatccccctttcgccagctggcgtaatagcgaagaggcccgcaccgatcgcccttcccaacagttgcgcagcctgaatggcgaatgg
aaattgtaagcgttaatattttgttaaaattcgcgttaaatttttgttaaatcagctcattttttaaccaataggccgaaatcggcaaaatcccttataaatcaaaag
aatagaccgagatagggttgagtgttgttccagtttggaacaagagtccactattaaagaacgtggactccaacgtcaaagggcgaaaaaccgtctatcag
ggcgatggcccactacgtgaaccatcaccctaatcaagttttttggggtcgaggtgccgtaaagcactaaatcggaaccctaaagggagcccccgatttag
agcttgacggggaaagccggcgaacgtggcgagaaaggaagggaagaaagcgaaaggagcgggcgctagggcgctggcaagtgtagcggtcac
gctgcgcgtaaccaccacacccgccgcgcttaatgcgccgctacagggcgcgtcaggtggcacttttcggggaaatgtgcgcggaacccctatttgtttatttt
tctaaatacattcaaatatgtatccgctcatgagacaataaccctgataaatgcttcaataatattgaaaaaggaagagtatgagtattcaacatttccgtgtcg
cccttattcccttttttgcggcattttgccttcctgtttttgctcacccagaaacgctggtgaaagtaaaagatgctgaagatcagttgggtgcacgagtgggttaca
tcgaactggatctcaacagcggtaagatccttgagagttttcgccccgaagaacgttttccaatgatgagcacttttaaagttctgctatgtggcgcggtattatc
ccgtattgacgccgggcaagagcaactcggtcgccgcatacactattctcagaatgacttggttgagtactcaccagtcacagaaaagcatcttacggatgg
catgacagtaagagaattatgcagtgctgccataaccatgagtgataacactgcggccaacttacttctgacaacgatcggaggaccgaaggagctaacc
gcttttttgcacaacatgggggatcatgtaactcgccttgatcgttgggaaccggagctgaatgaagccataccaaacgacgagcgtgacaccacgatgcc
tgtagcaatggcaacaacgttgcgcaaactattaactggcgaactacttactctagcttcccggcaacaattaatagactggatggaggcggataaagttgc
aggaccacttctgcgctcggcccttccggctggctggtttattgctgataaatctggagccggtgagcgtgggtctcgcggtatcattgcagcactggggccag

atggtaagccctcccgtatcgtagttatctacacgacggggagtcaggcaactatggatgaacgaaatagacagatcgctgagataggtgcctcactgatta
agcattggtaactgtcagaccaagtttactcatatatactttagattgatttaaaacttcatttttaatttaaaaggatctaggtgaagatcctttttgataatctcatga
ccaaaatcccttaacgtgagttttcgttccactgagcgtcagaccccgtagaaaagatcaaaggatcttcttgagatcctttttttctgcgcgtaatctgctgcttgc
aaacaaaaaaaccaccgctaccagcggtggtttgtttgccggatcaagagctaccaactctttttccgaaggtaactggcttcagcagagcgcagatacca
aatactgttcttctagtgtagccgtagttaggccaccacttcaagaactctgtagcaccgcctacatacctcgctctgctaatcctgttaccagtggctgctgcca
gtggcgataagtcgtgtcttaccgggttggactcaagacgatagttaccggataaggcgcagcggtcgggctgaacggggggttcgtgcacacagcccag
cttggagcgaacgacctacaccgaactgagatacctacagcgtgagctatgagaaagcgccacgcttcccgaagggagaaaggcggacaggtatccg
gtaagcggcagggtcggaacaggagagcgcacgagggagcttccagggggaaacgcctggtatctttatagtcctgtcgggtttcgccacctctgacttga
gcgtcgatttttgtgatgctcgtcaggggggcggagcctatggaaaaacgccagcaacgcggcctttttacggttcctggccttttgctggccttttgctcacatgt
tctttcctgcgttatcccctgattctgtggataaccgtattaccgcctttgagtgagctgataccgctcgccgcagccgaacgaccgagcgcagcgagtcagtg
agcgaggaagcggaagagcgcccaatacgcaaaccgcctctccccgcgcgttggccgattcattaatgcagctggcacgacaggtttcccgactggaa
agcgggcagtgagcgcaacgcaattaatgtgagttagctcactcattaggcaccccaggctttacactttatgcttccggctcgtatgttgtgtggaattgtgag
cggataacaatttcacacaggaaacagctatgaccatgattacgccaagcgcgcaattaaccctcactaaagggaacaaaagctggagctccaccgcg
gtggcggccgcctccacctccccagcaacatcaggctgggtgccatcctgcacagccactgcttttcacttccaggcctttgctagtattgttttctatgcctaca
ggccgccaccaccccacccacttttcctgtgtttgccttcccctgccgacacccaagcttgaagacctgatccaggctgggttaggcagcccttctttgtgttctg
tgatgagcggcgaactaaaacgcttgagatcgatgattaagggatctgtagggcgcagtagtccagggtttccttgatgatgtcatacttatcctgtccctttttttt
tccacagactcccaagaagagtcaccagaaatctctgagcttttctaaaactacaccaagaaggatctctcatacaccacaaactccgttgtatactccaga
aaggctgcagaagtcccctgcaaaaatgacccctacaaagcaggcagcttttaaggagtccttaaaagactcctcctcacccggccatgactcaccattgg
attcaaaaatcactcctcaaaaacgacatacccaggcaggagaaggtacctctcttgaaacgaagacaccaagaactcctaagaggcaaggtactcag
ccgcctgggtttttgccaaactgtacttggccacattcagtgaattccagtccagaaagcccctcctgtccagcccctccaacttcatcgactgcccagcccag
gagagagtgtctcactcccatcagagaccctctcagaacacctccgagagcagcagccttcatgggcacgcctcagaatcaaacacaccaacagcccc
atgtcctcagagctgctcgggcagaggaaccagcccagaaactaaaggataaagctatcaaaactccaaaaagaccagggaattcaactgtgacttctt
ccccacccgtgacccccaaaaagctcttcacctctcctttatgtgatgtctccaagaagagtccatttaggaaatctaaaatagagtgtccttccccaggagaa
ctggatcagaaagagccccagatgtcacccagcgtagctgcatctctctcctgccctgttccctcaactcccccctgaactctcacagagagctacattggaca
ccgtccctcctccaccccccttctaaagttgggaaacggtgtagaaagacctctgatcccagaaggagcatcgtggagtgtcagcctgatgcctccgctactc
ctggggttggcacagctgacagcccagctgcccccacagactctagggatgaccagaagggactgagcctctctcctcagtatcctcctgaaagacgggg
ctacccaggccccggtctcaggagtgattggcatgcatcctctcctctgctcattacaagtgacacagagcatgtcactctcctcagtgaagccgaacaccat
ggcattggtgacttgaaaagtaacgtcttatcagtggaagagggtgaggggctaaggacagcagatgctgagaagtcttctctgtctcaccccgggattccc
ccatctcctccttcctgtgggcctggctctcctctgatgccttcctgtgacgtgcactgtaccacagatgggagacagtgccaggcttcggcacaactagacaa
cctgccagcatcagcttggcattccacagactctgccagcccacagacctatgaggttgagctggagatgcaagcttctggccttcccaaacttcgaattaag
aagatagaccccagctcttcattagaggctgagcccctcagcaaggaggagagctctctgggagaagagagcttcctccctgctctcagcatgcccgggg
ccagcaggtccttaagcaaacctgaacccacctatgtgtcacccccctgcccccgcctctcccacagcacacccggcaagagcagggggcaaacctac
atctgccaggcctgtacccccacccacggcccttctagtacccccctctccatttcaaacagatggggttccttggacaccatcccccaagcacagtgggaag
acaactccagacataattaaagactggcccaggaggaagagggcggtgggctgtggcgccggctcctcttccgggaggggcgaggtcggtgcagacct
tcccgggagcctgtcactgcttgagtcagagggcaaggaccacggccttgaactcagcatccacaggacgcccatcttggaggattttgagctcgaggga
gtgtgccagctcccagaccagtcgcctcccaggaacagcatgcctaaggccgaggaagcctcttcctggggacagtttgggttgagttccgggaagagag
tcctgttggccaaggaagaagctgaccgtggagccaaaaggatctgtgatcttcgcgaggactccgaggtgagtaagagtaaagaggggtctccaagttg
gagtgcatggcagctaccctccacgggagacgaagaggtgtttgtttccggctccaccccacctcccagctgtgccgtgcggagctgcctctctgccagtgc
cctccaggctctgacccagtctccgctgctgttccaggggaaaacaccttcctctcagagcaaagaccccagagatgaggatgtggatgttcttccctccact
gtagaagactctcctttcagtcgcgctttctccaggaggcgccccatcagcagaacttatacacggaagaagctcatgggaacctggctggaggacggcg
gtggctctggaggtggtggatcgggaggtggctctatggtgagtaagggcgaggagcttttcaccggagtagtacccatcttggtcgagttggacggtgacg
taaacggtcacaagttcagtgttcgcggtgaaggtgaaggcgatgctaccaacggcaagctgaccctgaagttcatctgcaccaccggtaagcttcctgtac
cttggcctaccttggtgaccaccttcggttacggtgtggcttgcttcagtcgctaccctgatcacatgaagcagcacgacttcttcaagtcagctatgcccgaag
gttacgttcaggagcgcactatctccttcaaggacgacggtacctacaaaactcgcgctgaggtaaagttcgagggtgacaccttggtgaaccgcatcgag
ctgaagggcatcgacttcaaggaggacggcaacatccttgggcacaagctggagtacaacttcaacagccacaacgtctatatcacggctgacaagcag
aagaacggcatcaaggctaacttcaagatccgccacaacgttgaggacggtagtgtgcagttggctgaccactaccagcagaacactcccatcggtgatg
gtcccgtattgctccccgacaaccactacctgagccatcagtccaagctgagcaaagaccccaacgagaaacgcgatcacatggtcctgctggagttcgt
aaccgctgctggaattacacatggcatggacgagctgtacaaggactacaaggaccatgacggcgactataaggaccatgacatcgactacaaggacg
acgatgacaagtaatctagataactgatcataatcagccataccacatttgtagaggttttacttgctttaaaaaacctcccacacctccccctgaacctgaaa
cataaaatgaatgcaattgttgttgttaacttgtttattgcagcttataatggttacaaataaagcaatagcatcacaaatttcacaaataaagcatttttttcactgc
attctagttgtggtttgtccaaactcatcaatgtatcttaacgcgtcgatcatattcaataacccttaatataacttcgtataatgtatgctatacgaagttattaggtct
gaagaggagtttacgtccagccaagcttaggatctcgacctcgaaattctaccgggtaggggaggcgcttttcccaaggcagtctggagcatgcgctttagc

agccccgctgggcacttggcgctacacaagtggcctctggcctcgcacacattccacatccaccggtaggcgccaaccgactccgttctttggtggcccctt
cgcgccaccttctactcctcccctagtcaggaagttcccccccgccccgcagctcgcgtcgtgcaggacgtgacaaatggaagtagcacgtctcactagtct
cgtgcagatggacagcaccgctgagcaatggaagcgggtaggcctttggggcagcggccaatagcagctttgctccttcgctttctgggctcagaggctgg
gaaggggtgggtccgggggcgggctcaggggcgggctcaggggcggggcgggcgcccgaaggtcctccggaggcccggcattctgcacgcttcaaa
agcgcacgtctgccgcgctgttctcctcttcctcatctccgggcctttcgacctgcatccatctagatctcgagcagctgaagcttaccgctagcatggatagatc
cggaaagcctgaactcaccgcgacgtctgtcgagaagtttctgatcgaaaagttcgacagcgtctccgacctgatgcagctctcggagggcgaagaatctc
gtgctttcagcttcgatgtaggagggcgtggat

>pBlueScriptIISKPlus_TICRRIntronTargeting3_TICRR_8A (9C1)
atgtcctgcgggtaaatagctgcgccgatggtttctacaaagatcgttatgtttatcggcactttgcatcggccgcgctcccgattccggaagtgcttgacattgg
ggaattcagcgagagcctgacctattgcatctcccgccgtgcacagggtgtcacgttgcaagacctgcctgaaaccgaactgcccgctgttctgcagccggt
cgcggaggccatggatgcgatcgctgcggccgatcttagccagacgagcgggttcggcccattcggaccgcaaggaatcggtcaatacactacatggcg
tgatttcatatgcgcgattgctgatccccatgtgtatcactggcaaactgtgatggacgacaccgtcagtgcgtccgtcgcgcaggctctcgatgagctgatgct
ttgggccgaggactgccccgaagtccggcacctcgtgcacgcggatttcggctccaacaatgtcctgacggacaatggccgcataacagcggtcattgact
ggagcgaggcgatgttcggggattcccaatacgaggtcgccaacatcttcttctggaggccgtggttggcttgtatggagcagcagacgcgctacttcgagc
ggaggcatccggagcttgcaggatcgccgcggctccgggcgtatatgctccgcattggtcttgaccaactctatcagagcttggttgacggcaatttcgatgat
gcagcttgggcgcagggtcgatgcgacgcaatcgtccgatccggagccgggactgtcgggcgtacacaaatcgcccgcagaagcgcggccgtctgga
ccgatggctgtgtagaagtactcgccgatagtggaaaccgacgccccagcactcgtccgagggcaaaggaataggctagccgcccgccccacgaccc
gcagcgcccgaccgaaaggagcgcacgaccccatgcatcgatgatatcagatccccgggatgcagaaattgatgatctattaaacaataaagatgtcca
ctaaaatggaagtttttcctgtcatactttgttaagaagggtgagaacagagtacctacattttgaatggaaggattggagctacgggggtgggggtggggtg
ggattagataaatgcctgctctttactgaaggctctttactattgctttatgataatgtttcatagttggatatcataatttaaacaagcaaaaccaaattaagggcc
agctcattcctcccactcatgatctatagatctatagatctctcgtgggatcattgtttttctcttgattcccactttgtggttctaagtactgtggtttccaaatgtgtcag
tttcatagcctgaagaacgagatcagcagcctctgttccacatacacttcattctcagtattgttttgccaagttctaattccatcagaagctggtcgagatccgga
acccttaatataacttcgtataatgtatgctatacgaagttattaggtccctcgaagaggttcactaggatccgcatgtccctgtcactgcaccaatcacattgtat
tttcacctgcgtatgccccactgaacaggaaatgccagctagactgagagtcccctgacgggatgaaggaaggctgtaggaaagataagcattgatttcctt
acctgttggagcttttcactacgtatttttgcaagaaggcagggaaatttttattggggagtgtcttttctgaatcaagcttatcgataccgtcgacctcgaggggg
ggcccggtacccaattcgccctatagtgagtcgtattacgcgcgctcactggccgtcgttttacaacgtcgtgactgggaaaaccctggcgttacccaacttaa
tcgccttgcagcacatccccctttcgccagctggcgtaatagcgaagaggcccgcaccgatcgcccttcccaacagttgcgcagcctgaatggcgaatgg
aaattgtaagcgttaatattttgttaaaattcgcgttaaatttttgttaaatcagctcattttttaaccaataggccgaaatcggcaaaatcccttataaatcaaaag
aatagaccgagatagggttgagtgttgttccagtttggaacaagagtccactattaaagaacgtggactccaacgtcaaagggcgaaaaaccgtctatcag
ggcgatggcccactacgtgaaccatcaccctaatcaagttttttggggtcgaggtgccgtaaagcactaaatcggaaccctaaagggagcccccgatttag
agcttgacggggaaagccggcgaacgtggcgagaaaggaagggaagaaagcgaaaggagcgggcgctagggcgctggcaagtgtagcggtcac
gctgcgcgtaaccaccacacccgccgcgcttaatgcgccgctacagggcgcgtcaggtggcacttttcggggaaatgtgcgcggaacccctatttgtttatttt
tctaaatacattcaaatatgtatccgctcatgagacaataaccctgataaatgcttcaataatattgaaaaaggaagagtatgagtattcaacatttccgtgtcg
cccttattcccttttttgcggcattttgccttcctgtttttgctcacccagaaacgctggtgaaagtaaaagatgctgaagatcagttgggtgcacgagtgggttaca
tcgaactggatctcaacagcggtaagatccttgagagttttcgccccgaagaacgttttccaatgatgagcacttttaaagttctgctatgtggcgcggtattatc
ccgtattgacgccgggcaagagcaactcggtcgccgcatacactattctcagaatgacttggttgagtactcaccagtcacagaaaagcatcttacggatgg
catgacagtaagagaattatgcagtgctgccataaccatgagtgataacactgcggccaacttacttctgacaacgatcggaggaccgaaggagctaacc
gcttttttgcacaacatgggggatcatgtaactcgccttgatcgttgggaaccggagctgaatgaagccataccaaacgacgagcgtgacaccacgatgcc
tgtagcaatggcaacaacgttgcgcaaactattaactggcgaactacttactctagcttcccggcaacaattaatagactggatggaggcggataaagttgc
aggaccacttctgcgctcggcccttccggctggctggtttattgctgataaatctggagccggtgagcgtgggtctcgcggtatcattgcagcactggggccag
atggtaagccctcccgtatcgtagttatctacacgacggggagtcaggcaactatggatgaacgaaatagacagatcgctgagataggtgcctcactgatta
agcattggtaactgtcagaccaagtttactcatatatactttagattgatttaaaacttcatttttaatttaaaaggatctaggtgaagatcctttttgataatctcatga
ccaaaatcccttaacgtgagttttcgttccactgagcgtcagaccccgtagaaaagatcaaaggatcttcttgagatcctttttttctgcgcgtaatctgctgcttgc
aaacaaaaaaaccaccgctaccagcggtggtttgtttgccggatcaagagctaccaactctttttccgaaggtaactggcttcagcagagcgcagatacca
aatactgttcttctagtgtagccgtagttaggccaccacttcaagaactctgtagcaccgcctacatacctcgctctgctaatcctgttaccagtggctgctgcca
gtggcgataagtcgtgtcttaccgggttggactcaagacgatagttaccggataaggcgcagcggtcgggctgaacggggggttcgtgcacacagcccag
cttggagcgaacgacctacaccgaactgagatacctacagcgtgagctatgagaaagcgccacgcttcccgaagggagaaaggcggacaggtatccg
gtaagcggcagggtcggaacaggagagcgcacgagggagcttccagggggaaacgcctggtatctttatagtcctgtcgggtttcgccacctctgacttga
gcgtcgatttttgtgatgctcgtcaggggggcggagcctatggaaaaacgccagcaacgcggcctttttacggttcctggccttttgctggccttttgctcacatgt
tctttcctgcgttatcccctgattctgtggataaccgtattaccgcctttgagtgagctgataccgctcgccgcagccgaacgaccgagcgcagcgagtcagtg
agcgaggaagcggaagagcgcccaatacgcaaaccgcctctccccgcgcgttggccgattcattaatgcagctggcacgacaggtttcccgactggaa

agcgggcagtgagcgcaacgcaattaatgtgagttagctcactcattaggcaccccaggctttacactttatgcttccggctcgtatgttgtgtggaattgtgag
cggataacaatttcacacaggaaacagctatgaccatgattacgccaagcgcgcaattaaccctcactaaagggaacaaaagctggagctccaccgcg
gtggcggccgcctccacctccccagcaacatcaggctgggtgccatcctgcacagccactgcttttcacttccaggcctttgctagtattgttttctatgcctaca
ggccgccaccaccccacccacttttcctgtgtttgccttcccctgccgacacccaagcttgaagacctgatccaggctgggttaggcagcccttctttgtgttctg
tgatgagcggcgaactaaaacgcttgagatcgatgattaagggatctgtagggcgcagtagtccagggtttccttgatgatgtcatacttatcctgtccctttttttt
tccacagactcccaagaagagtcaccagaaatctctgagcttttctaaaactacaccaagaaggatctctcatacaccacaaactccgttgtatactccaga
aaggctgcagaagtcccctgcaaaaatgaccccctacaaagcaggcagcttttaaggagtccttaaaagactcctcctcacccggccatgactcaccattgg
attcaaaaatcactcctcaaaaacgacatacccaggcaggagaaggtacctctcttgaaacgaagacaccaagaactcctaagaggcaaggtactcag
ccgcctgggtttttgccaaactgtacttggccacattcagtgaattccagtccagaaagcccctcctgtccagcccctccaacttcatcgactgcccagcccag
gagagagtgtctcactcccatcagagaccctctcagaacacctccgagagcagcagccttcatgggcacgcctcagaatcaaacacaccaacagcccc
atgtcctcagagctgctcgggcagaggaaccagcccagaaactaaaggataaagctatcaaaactccaaaaagaccggggaattcaactgtgacttctt
ccccacccgtgacccccaaaaagctcttcacctctcctttatgtgatgtctccaagaagagtccatttaggaaatctaaaatagagtgtccttccccaggagaa
ctggatcagaaagagccccagatgtcacccagcgtagctgcatctctctcctgccctgttccctcaactcccccctgaactctcacagagagctacattggaca
ccgtccctcctccaccccccttctaaagttgggaaacggtgtagaaagacctctgatcccagaaggagcatcgtggagtgtcagcctgatgcctccgctactc
ctggggttggcacagctgacagcccagctgcccccacagactctagggatgaccagaagggactgagcctctctcctcagtatcctcctgaaagacgggg
ctacccagtgccccggtctcaggagtgattggcatgcatcctctcctctgctcattacaagtgacacagagcatgtcactctcctcagtgaagccgaacacca
tggcattggtgacttgaaaagtaacgtcttatcagtggaagagggtgaggggctaaggacagcagatgctgagaagtcttctctgtctcaccccgggattcc
cccatctcctccttcctgtgggcctggctctcctctgatgccttcctgtgacgtgcactgtaccacagatgggagacagtgccaggcttcggcacaactagaca
acctgccagcatcagcttggcattccacagactctgctagcccacagacctatgcggttgcgctggcgatgcaagcttctggccttcccgcacttgcaattgc
ggcgatagcccccagctcttcattagaggccgagccactcagcaaggaggaatcctctctgggtgaggagtcatttctgccagcactgtccatgccgaggg
ctagcaggtccttaagcaaacctgaacccacctatgtgtcacccccctgcccccgcctctcccacagcacacctggcaagagcagggggcaaacctaca
tctgccaggcctgtacccccacccacggcccttctagtacccccctctccatttcaaacagatggggttccttggacaccatcccccaagcacagtgggaaga
caactccagacataattaaagactggcccaggaggaagagggcggtgggctgtggcgccggctcctcttccgggaggggcgaggtcggtgcagacctt
cccgggagcctgtcactgcttgagtcagagggcaaggaccacggccttgaactcagcatccacaggacgcccatcttggaggattttgagctcgagggag
tgtgccagctcccagaccagtcgcctcccaggaacagcatgcctaaggccgaggaagcctcttcctggggacagtttgggttgagttccaggaagagagt
cctgttggccaaggaagaagctgaccgtggagccaaaaggatctgtgatcttcgcgaggactccgaggtgagtaagagtaaagaggggtctccaagttg
gagtgcatggcagctaccctccacgggagacgaagaggtgtttgtttccggctccaccccacctcccagctgtgccgtgcggagctgcctctctgccagtgc
cctccaggctctgacccagtctccgctgctgttccaggggaaaacaccttcctctcagagcaaagaccccagagatgaggatgtggatgttcttccctccact
gtagaagactctcctttcagtcgcgctttctccaggaggcgccccatcagcagaacttatacacggaagaagctcatgggaacctggctggaggacggcg
gtggctctggaggtggtggatcgggaggtggctctatggtgagtaagggcgaggagcttttcaccggagtagtacccatcttggtcgagttggacggtgacg
taaacggtcacaagttcagtgttcgcggtgaaggtgaaggcgatgctaccaacggcaagctgaccctgaagttcatctgcaccaccggtaagcttcctgtac
cttggcctaccttggtgaccaccttcggttacggtgtggcttgcttcagtcgctaccctgatcacatgaagcagcacgacttcttcaagtcagctatgcccgaag
gttacgttcaggagcgcactatctccttcaaggacgacggtacctacaaaactcgcgctgaggtaaagttcgagggtgacaccttggtgaaccgcatcgag
ctgaagggcatcgacttcaaggaggacggcaacatccttgggcacaagctggagtacaacttcaacagccacaacgtctatatcacggctgacaagcag
aagaacggcatcaaggctaacttcaagatccgccacaacgttgaggacggtagtgtgcagttggctgaccactaccagcagaacactcccatcggtgatg
gtcccgtattgctccccgacaaccactacctgagccatcagtccaagctgagcaaagaccccaacgagaaacgcgatcacatggtcctgctggagttcgt
aaccgctgctggaattacacatggcatggacgagctgtacaaggactacaaggaccatgacggcgactataaggaccatgacatcgactacaaggacg
acgatgacaagtaatctagataactgatcataatcagccataccacatttgtagaggttttacttgctttaaaaaacctcccacacctccccctgaacctgaaa
cataaaatgaatgcaattgttgttgttaacttgtttattgcagcttataatggttacaaataaagcaatagcatcacaaatttcacaaataaagcatttttttcactgc
attctagttgtggtttgtccaaactcatcaatgtatcttaacgcgtcgatcatattcaataacccttaatataacttcgtataatgtatgctatacgaagttattaggtct
gaagaggagtttacgtccagccaagcttaggatctcgacctcgaaattctaccgggtaggggaggcgcttttcccaaggcagtctggagcatgcgctttagc
agccccgctgggcacttggcgctacacaagtggcctctggcctcgcacacattccacatccaccggtaggcgccaaccgactccgttctttggtggcccctt
cgcgccaccttctactcctcccctagtcaggaagttcccccccgccccgcagctcgcgtcgtgcaggacgtgacaaatggaagtagcacgtctcactagtct
cgtgcagatggacagcaccgctgagcaatggaagcgggtaggcctttggggcagcggccaatagcagctttgctccttcgctttctgggctcagaggctgg
gaaggggtgggtccgggggcgggctcaggggcgggctcaggggcggggcgggcgcccgaaggtcctccggaggcccggcattctgcacgcttcaaa
agcgcacgtctgccgcgctgttctcctcttcctcatctccgggcctttcgacctgcatccatctagatctcgagcagctgaagcttaccgctagcatggatagatc
cggaaagcctgaactcaccgcgacgtctgtcgagaagtttctgatcgaaaagttcgacagcgtctccgacctgatgcagctctcggagggcgaagaatctc
gtgctttcagcttcgatgtaggagggcgtggat

>pBlueScriptIISKPlus_TICRRIntronTargeting3_TICRR_ΔSBI (9B9)
ggtaaatagctgcgccgatggtttctacaaagatcgttatgtttatcggcactttgcatcggccgcgctcccgattccggaagtgcttgacattggggaattcag
cgagagcctgacctattgcatctcccgccgtgcacagggtgtcacgttgcaagacctgcctgaaaccgaactgcccgctgttctgcagccggtcgcggagg

ccatggatgcgatcgctgcggccgatcttagccagacgagcgggttcggcccattcggaccgcaaggaatcggtcaatacactacatggcgtgatttcatat
gcgcgattgctgatccccatgtgtatcactggcaaactgtgatggacgacaccgtcagtgcgtccgtcgcgcaggctctcgatgagctgatgctttgggccga
ggactgccccgaagtccggcacctcgtgcacgcggatttcggctccaacaatgtcctgacggacaatggccgcataacagcggtcattgactggagcga
ggcgatgttcggggattcccaatacgaggtcgccaacatcttcttctggaggccgtggttggcttgtatggagcagcagacgcgctacttcgagcggaggca
tccggagcttgcaggatcgccgcggctccgggcgtatatgctccgcattggtcttgaccaactctatcagagcttggttgacggcaatttcgatgatgcagcttg
ggcgcagggtcgatgcgacgcaatcgtccgatccggagccgggactgtcgggcgtacacaaatcgcccgcagaagcgcggccgtctggaccgatggc
tgtgtagaagtactcgccgatagtggaaaccgacgccccagcactcgtccgagggcaaaggaataggctagccgcccgccccacgacccgcagcgcc
cgaccgaaaggagcgcacgaccccatgcatcgatgatatcagatccccgggatgcagaaattgatgatctattaaacaataaagatgtccactaaaatgg
aagtttttcctgtcatactttgttaagaagggtgagaacagagtacctacattttgaatggaaggattggagctacgggggtgggggtggggtgggattagata
aatgcctgctctttactgaaggctctttactattgctttatgataatgtttcatagttggatatcataatttaaacaagcaaaaccaaattaagggccagctcattcct
cccactcatgatctatagatctatagatctctcgtgggatcattgtttttctcttgattcccactttgtggttctaagtactgtggtttccaaatgtgtcagtttcatagcct
gaagaacgagatcagcagcctctgttccacatacacttcattctcagtattgttttgccaagttctaattccatcagaagctggtcgagatccggaacccttaata
taacttcgtataatgtatgctatacgaagttattaggtccctcgaagaggttcactaggatccgcatgtccctgtcactgcaccaatcacattgtattttcacctgcg
tatgccccactgaacaggaaatgccagctagactgagagtcccctgacgggatgaaggaaggctgtaggaaagataagcattgatttccttacctgttgga
gcttttcactacgtatttttgcaagaaggcagggaaatttttattggggagtgtcttttctgaatcaagcttatcgataccgtcgacctcgagggggggcccggta
cccaattcgccctatagtgagtcgtattacgcgcgctcactggccgtcgttttacaacgtcgtgactgggaaaaccctggcgttacccaacttaatcgccttgca
gcacatccccctttcgccagctggcgtaatagcgaagaggcccgcaccgatcgcccttcccaacagttgcgcagcctgaatggcgaatggaaattgtaag
cgttaatattttgttaaaattcgcgttaaatttttgttaaatcagctcattttttaaccaataggccgaaatcggcaaaatcccttataaatcaaaagaatagaccg
agatagggttgagtgttgttccagtttggaacaagagtccactattaaagaacgtggactccaacgtcaaagggcgaaaaaccgtctatcagggcgatggc
ccactacgtgaaccatcaccctaatcaagttttttggggtcgaggtgccgtaaagcactaaatcggaaccctaaagggagcccccgatttagagcttgacgg
ggaaagccggcgaacgtggcgagaaaggaagggaagaaagcgaaaggagcgggcgctagggcgctggcaagtgtagcggtcacgctgcgcgtaa
ccaccacacccgccgcgcttaatgcgccgctacagggcgcgtcaggtggcacttttcggggaaatgtgcgcggaacccctatttgtttatttttctaaatacatt
caaatatgtatccgctcatgagacaataaccctgataaatgcttcaataatattgaaaaaggaagagtatgagtattcaacatttccgtgtcgcccttattcccttt
tttgcggcattttgccttcctgtttttgctcacccagaaacgctggtgaaagtaaaagatgctgaagatcagttgggtgcacgagtgggttacatcgaactggat
ctcaacagcggtaagatccttgagagttttcgccccgaagaacgttttccaatgatgagcacttttaaagttctgctatgtggcgcggtattatcccgtattgacg
ccgggcaagagcaactcggtcgccgcatacactattctcagaatgacttggttgagtactcaccagtcacagaaaagcatcttacggatggcatgacagta
agagaattatgcagtgctgccataaccatgagtgataacactgcggccaacttacttctgacaacgatcggaggaccgaaggagctaaccgcttttttgcac
aacatgggggatcatgtaactcgccttgatcgttgggaaccggagctgaatgaagccataccaaacgacgagcgtgacaccacgatgcctgtagcaatg
gcaacaacgttgcgcaaactattaactggcgaactacttactctagcttcccggcaacaattaatagactggatggaggcggataaagttgcaggaccactt
ctgcgctcggcccttccggctggctggtttattgctgataaatctggagccggtgagcgtgggtctcgcggtatcattgcagcactggggccagatggtaagcc
ctcccgtatcgtagttatctacacgacggggagtcaggcaactatggatgaacgaaatagacagatcgctgagataggtgcctcactgattaagcattggta
actgtcagaccaagtttactcatatatactttagattgatttaaaacttcatttttaatttaaaaggatctaggtgaagatcctttttgataatctcatgaccaaaatcc
cttaacgtgagttttcgttccactgagcgtcagaccccgtagaaaagatcaaaggatcttcttgagatcctttttttctgcgcgtaatctgctgcttgcaaacaaaa
aaaccaccgctaccagcggtggtttgtttgccggatcaagagctaccaactctttttccgaaggtaactggcttcagcagagcgcagataccaaatactgttct
tctagtgtagccgtagttaggccaccacttcaagaactctgtagcaccgcctacatacctcgctctgctaatcctgttaccagtggctgctgccagtggcgataa
gtcgtgtcttaccgggttggactcaagacgatagttaccggataaggcgcagcggtcgggctgaacggggggttcgtgcacacagcccagcttggagcga
acgacctacaccgaactgagatacctacagcgtgagctatgagaaagcgccacgcttcccgaagggagaaaggcggacaggtatccggtaagcggc
agggtcggaacaggagagcgcacgagggagcttccagggggaaacgcctggtatctttatagtcctgtcgggtttcgccacctctgacttgagcgtcgatttt
tgtgatgctcgtcaggggggcggagcctatggaaaaacgccagcaacgcggcctttttacggttcctggccttttgctggccttttgctcacatgttctttcctgcg
ttatcccctgattctgtggataaccgtattaccgcctttgagtgagctgataccgctcgccgcagccgaacgaccgagcgcagcgagtcagtgagcgagga
agcggaagagcgcccaatacgcaaaccgcctctccccgcgcgttggccgattcattaatgcagctggcacgacaggtttcccgactggaaagcgggcag
tgagcgcaacgcaattaatgtgagttagctcactcattaggcaccccaggctttacactttatgcttccggctcgtatgttgtgtggaattgtgagcggataacaa
tttcacacaggaaacagctatgaccatgattacgccaagcgcgcaattaaccctcactaaagggaacaaaagctggagctccaccgcggtggcggccg
cctccacctccccagcaacatcaggctgggtgccatcctgcacagccactgcttttcacttccaggcctttgctagtattgttttctatgcctacaggccgccacc
accccacccacttttcctgtgtttgccttcccctgccgacacccaagcttgaagacctgatccaggctgggttaggcagcccttctttgtgttctgtgatgagcgg
cgaactaaaacgcttgagatcgatgattaagggatctgtagggcgcagtagtccagggtttccttgatgatgtcatacttatcctgtcccttttttttccacagactc
ccaagaagagtcaccagaaatctctgagcttttctaaaactacaccaagaaggatctctcatacaccacaaactccgttgtatactccagaaaggctgcag
aagtcccctgcaaaaatgacccctacaaagcaggcagcttttaaggagtccttaaaagactcctcctcacccggccatgactcaccattggattcaaaaatc
actcctcaaaaacgacatacccaggcaggagaaggtacctctcttgaaacgaagacaccaagaactcctaagaggcaaggtactcagccgcctgggttt
ttgccaaactgtacttggccacattcagtgaattccagtccagaaagcccctcctgtccagcccctccaacttcatcgactgcccagcccaggagagagtgtc
tcactcccatcagagaccctctcagaacacctccgagagcagcagccttcatgggcacgcctcagaatcaaacacaccaacagccccatgtcctcagag
ctgctcgggcagaggaaccagcccagaaactaaaggataaagctatcaaaactccaaaaagaccagggaattcaactgtgacttcttccccacccgtga

cccccaaaaagctcttcacctctcctttatgtgatgtctccaagaagagtccatttaggaaatctaaaatagagtgtccttccccaggagaactggatcagaaa
gagccccagatgtcacccagcgtagctgcatctctctcctgccctgttccctcaactccccctgaactctcacagagagctacattggacaccgtccctcctcc
accccccttctaaagttgggaaacggtgtagaaagacctctgatcccagaaggagcatcgtggagtgtcagcctgatgcctccgctactcctggggttggcac
agctgacagcccagctgcccccacagactctagggatgaccagaagggactgagcctctctcctcagtatcctcctgaaagacggggctacccaggccc
cggtctcaggagtgattggcatgcatcctctcctctgctcattacaagtgacacagagcatgtcactctcctcagtgaagccgaacaccatggcattggtgact
tgaaaagtaacgtcttatcacaaacagatggggttccttggacaccatcccccaagcacagtgggaagacaactccagacataattaaagactggccca
ggaggaagagggcggtgggctgtggcgccggctcctcttccgggaggggcgaggtcggtgcagaccttcccgggagcctgtcactgcttgagtcagagg
gcaaggaccacggccttgaactcagcatccacaggacgcccatcttggaggattttgagctcgagggagtgtgccagctcccagaccagtcgcctcccag
gaacagcatgcctaaggccgaggaagcctcttcctggggacagtttgggttgagttccaggaagagagtcctgttggccaaggaagaagctgaccgtgg
agccaaaaggatctgtgatcttcgcgaggactccgaggtgagtaagagtaaagaggggtctccaagttggagtgcatggcagctaccctccacgggaga
cgaagaggtgtttgtttccggctccaccccacctcccagctgtgccgtgcggagctgcctctctgccagtgccctccaggctctgacccagtctccgctgctgtt
ccaggggaaaacaccttcctctcagagcaaagaccccagagatgaggatgtggatgttcttccctccactgtagaagactctcctttcagtcgcgctttctcca
ggaggcgccccatcagcagaacttatacacggaagaagctcatgggaacctggctggaggacggcggtggctctggaggtggtggatcgggaggtgg
ctctatggtgagtaagggcgaggagcttttcaccggagtagtacccatcttggtcgagttggacggtgacgtaaacggtcacaagttcagtgttcgcggtgaa
ggtgaaggcgatgctaccaacggcaagctgaccctgaagttcatctgcaccaccggtaagcttcctgtaccttggcctaccttggtgaccaccttcggttacg
gtgtggcttgcttcagtcgctaccctgatcacatgaagcagcacgacttcttcaagtcagctatgcccgaaggttacgttcaggagcgcactatctccttcaag
gacgacggtacctacaaaactcgcgctgaggtaaagttcgagggtgacaccttggtgaaccgcatcgagctgaagggcatcgacttcaaggaggacgg
caacatccttgggcacaagctggagtacaacttcaacagccacaacgtctatatcacggctgacaagcagaagaacggcatcaaggctaacttcaagat
ccgccacaacgttgaggacggtagtgtgcagttggctgaccactaccagcagaacactcccatcggtgatggtcccgtattgctccccgacaaccactacc
tgagccatcagtccaagctgagcaaagaccccaacgagaaacgcgatcacatggtcctgctggagttcgtaaccgctgctggaattacacatggcatgga
cgagctgtacaaggactacaaggaccatgacggcgactataaggaccatgacatcgactacaaggacgacgatgacaagtaatctagataactgatca
taatcagccataccacatttgtagaggttttacttgctttaaaaaacctcccacacctccccctgaacctgaaacataaaatgaatgcaattgttgttgttaacttg
tttattgcagcttataatggttacaaataaagcaatagcatcacaaatttcacaaataaagcatttttttcactgcattctagttgtggtttgtccaaactcatcaatgt
atcttaacgcgtcgatcatattcaataacccttaatataacttcgtataatgtatgctatacgaagttattaggtctgaagaggagtttacgtccagccaagcttag
gatctcgacctcgaaattctaccgggtaggggaggcgcttttcccaaggcagtctggagcatgcgctttagcagccccgctgggcacttggcgctacacaa
gtggcctctggcctcgcacacattccacatccaccggtaggcgccaaccgactccgttctttggtggccccttcgcgccaccttctactcctcccctagtcagg
aagttcccccccgccccgcagctcgcgtcgtgcaggacgtgacaaatggaagtagcacgtctcactagtctcgtgcagatggacagcaccgctgagcaat
ggaagcgggtaggcctttggggcagcggccaatagcagctttgctccttcgctttctgggctcagaggctgggaaggggtgggtccgggggcgggctcag
gggcgggctcaggggcggggcgggcgcccgaaggtcctccggaggcccggcattctgcacgcttcaaaagcgcacgtctgccgcgctgttctcctcttcc
tcatctccgggcctttcgacctgcatccatctagatctcgagcagctgaagcttaccgctagcatggatagatccggaaagcctgaactcaccgcgacgtctg
tcgagaagtttctgatcgaaaagttcgacagcgtctccgacctgatgcagctctcggagggcgaagaatctcgtgctttcagcttcgatgtaggagggcgtgg
atatgtcctgcg

>XLone-CDT2 (9D8)
gctcccagccgatgcccttgacgactttgaccttgatatgctgcctgctgacgctcttgacgattttgaccttgacatgctccccgggggcagcggcgccacca
acttcagcctgctgaagcaggccggcgacgtggaggagaaccccggccccatggccaagcctttgtctcaagaagaatccaccctcattgaaagagca
acggctacaatcaacagcatccccatctctgaagactacagcgtcgccagcgcagctctctctagcgacggccgcatcttcactggtgtcaatgtatatcattt
tactgggggaccttgtgcagaactcgtggtgctgggcactgctgctgctgcggcagctggcaacctgacttgtatcgtcgcgatcggaaatgagaacaggg
gcatcttgagcccctgcggacggtgccgacaggtgcttctcgatctgcatcctgggatcaaagccatagtgaaggacagtgatggacagccgacggcagtt
gggattcgtgaattgctgccctctggttatgtgtgggagggctaaatctccagaggatcataatcagccataccacatttgtagaggttttacttgctttaaaaaa
cctcccacacctccccctgaacctgaaacataaaatgaatgcaattgttgttgttaacttgtttattgcagcttataatggttacaaataaagcaatagcatcaca
aatttcacaaataaagcatttttttcactgctaaaagttttgttactttatagaagaaattttgagtttttgtttttttttaataaataaataaacataaataaattgtttgttg
aatttattattagtatgtaagtgtaaatataataaaacttaatatctattcaaattaataaataaacctcgatatacagaccgataaaacacatgcgtcaattttac
gcatgattatctttaacgtacgtcacaatatgattatctttctagggttaagtcgaccttggcgtaatcatggtcatagctgtttcctgtgtgaaattgttatccgctcac
aattccacacaacatacgagccggaagcataaagtgtaaagcctggggtgcctaatgagtgagctaactcacattaattgcgttgcgctcactgcccgctttc
cagtcgggaaacctgtcgtgccagctgcattaatgaatcggccaacgcgcggggagaggcggtttgcgtattgggcgctcttccgcttcctcgctcactgact
cgctgcgctcggtcgttcggctgcggcgagcggtatcagctcactcaaaggcggtaatacggttatccacagaatcaggggataacgcaggaaagaaca
tgtgagcaaaaggccagcaaaaggccaggaaccgtaaaaaggccgcgttgctggcgtttttccataggctccgcccccctgacgagcatcacaaaaatc
gacgctcaagtcagaggtggcgaaacccgacaggactataaagataccaggcgtttccccctggaagctccctcgtgcgctctcctgttccgaccctgccg
cttaccggatacctgtccgcctttctcccttcgggaagcgtggcgctttctcatagctcacgctgtaggtatctcagttcggtgtaggtcgttcgctccaagctggg
ctgtgtgcacgaaccccccgttcagcccgaccgctgcgccttatccggtaactatcgtcttgagtccaacccggtaagacacgacttatcgccactggcagc
agccactggtaacaggattagcagagcgaggtatgtaggcggtgctacagagttcttgaagtggtggcctaactacggctacactagaagaacagtatttg

gtatctgcgctctgctgaagccagttaccttcggaaaaagagttggtagctcttgatccggcaaacaaaccaccgctggtagcggtggtttttttgtttgcaagc
agcagattacgcgcagaaaaaaaggatctcaagaagatcctttgatcttttctacggggtctgacgctcagtggaacgaaaactcacgttaagggattttggt
catgagattatcaaaaaggatcttcacctagatccttttaaattaaaaatgaagttttaaatcaatctaaagtatatatgagtaaacttggtctgacagttaccaat
gcttaatcagtgaggcacctatctcagcgatctgtctatttcgttcatccatagttgcctgactccccgtcgtgtagataactacgatacgggagggcttaccatct
ggccccagtgctgcaatgataccgcgagacccacgctcaccggctccagatttatcagcaataaaccagccagccggaagggccgagcgcagaagtg
gtcctgcaactttatccgcctccatccagtctattaattgttgccgggaagctagagtaagtagttcgccagttaatagtttgcgcaacgttgttgccattgctaca
ggcatcgtggtgtcacgctcgtcgtttggtatggcttcattcagctccggttcccaacgatcaaggcgagttacatgatcccccatgttgtgcaaaaaagcggtt
agctccttcggtcctccgatcgttgtcagaagtaagttggccgcagtgttatcactcatggttatggcagcactgcataattctcttactgtcatgccatccgtaag
atgcttttctgtgactggtgagtactcaaccaagtcattctgagaatagtgtatgcggcgaccgagttgctcttgcccggcgtcaatacgggataataccgcgc
cacatagcagaactttaaaagtgctcatcattggaaaacgttcttcggggcgaaaactctcaaggatcttaccgctgttgagatccagttcgatgtaacccact
cgtgcacccaactgatcttcagcatcttttactttcaccagcgtttctgggtgagcaaaaacaggaaggcaaaatgccgcaaaaaagggaataagggcga
cacggaaatgttgaatactcatactcttcctttttcaatattattgaagcatttatcagggttattgtctcatgagcggatacatatttgaatgtatttagaaaaataaa
caaataggggttccgcgcacatttccccgaaaagtgccacctgacgtctaagaaaccattattatcatgacattaacctataaaaataggcgtatcacgagg
ccctttcgtctcgcgcgtttcggtgatgacggtgaaaacctctgacacatgcagctcccggagacggtcacagcttgtctgtaagcggatgccgggagcaga
caagcccgtcagggcgcgtcagcgggtgttggcgggtgtcggggctggcttaactatgcggcatcagagcagattgtactgagagtgcaccatatgcggtg
tgaaataccgcacagatgcgtaaggagaaaataccgcatcaggcgccattcgccattcaggctgcgcaactgttgggaagggcgatcggtgcgggcctc
ttcgctattacgccagctggcgaaagggggatgtgctgcaaggcgattaagttgggtaacgccagggttttcccagtcacgacgttgtaaaacgacggcca
gtgaattcttaaccctagaaagatagtctgcgtaaaattgacgcatgcattcttgaaatattgctctctctttctaaatagcgcgaatccgtcgctgtgcatttagga
catctcagtcgccgcttggagctcccgtgaggcgtgcttgtcaatgcggtaagtgtcactgattttgaactataacgaccgcgtgagtcaaaatgacgcatgat
tatcttttacgtgacttttaagatttaactcatacgataattatattgttatttcatgttctacttacgtgataacttattatatatatatattttcttgttatagatatcaaacttgttt
attgcagcttataatggttacaaataaggcaatagcatcacaaatttcacaaataaggcatttttttcactgcattctagttttggtttgtccaaactcatcaatgtat
cttatcatgtctggatctcaaatccctcggaagctgcgcctgtcttaggttggagtgatacatttttatcacttttacccgtctttggattaggcagtagctctgacgg
ccctcctgtcttaggttagtgaaaaatgtcactctcttacccgtcattggctgtccagcttagctcgcaggggaggtggtctatcgaggctgatcagcgagcttct
agttacttgtcatcgtcatccttgtaatcgatgtcatgatctttataatcaccgtcatggtctttgtagtctgcggccgctaattctgttgagtgttcaggaccacagaa
gtcctcctgggactttctatggaagtatgtgcagattttcctcatggagctgggcgtgatggtgaccgggcttggcaatgtctttccgctctgtctcctggaattggg
tgtctgggatgacggacttcgtggagatggattctcagccttccgtttggctgccatggccaacaaccagtttttattctctggggaactattctctttgcctaccattt
cagacccttctccacaaggtctcaaaggaagaggtagcgttccacagctttctgaagcatacggactgataggagacggaggctctgagatactggtacc
agctccttcaattttgcttgatttggtaggacctagagagtccttactaaggtcttcctggttaccagcaaggcagcacagatccaaatgaagattttcaacttgg
ccatcaagctcagtcacacagttacaactcttcacacacttttgtttcacactctccagacagcttgagtctagcctcctctttactctatttctagactcagagcaa
gcctctgcttgggatgacttctggctcacaggaataagggcttttctcggagacatgatcttggtctccgaagcaggtggagtgatgggtggtgatgaggaag
gtgttcgggtcacccagtttctaatcgacatcttgaaagatgaaggtggcttgggagagacggaggagacagagcctcttctgttgatgggagaccgggcct
tggcaggagaggttttaatagagaacgtaggagtatttgaaggaagagggaggtctccagcacagcttggggcacaagctgcggatgacggggaagaa
ttggatggattgcactttaccctgggggctttggcaggagtactctggctactcgttactgttactaggccaggtcttgactctttttcttctgagaggcccaaccca
ccgtggaaagtttatcacctcctggtttctcctctaagcctctattcaagcgccagatttttagtgtattgtcatcagaacaggtagcaatctttgtgaagtcagatg
gacaccagcacacagacgtgacctcttgagaatgacccaggagcacagtaggaggttgccagggtgtggagaccttccatatgtaggcagcttcatcact
tgagccactgactaaaaactggtcatctggactaaggctggattttacataaaaggtagagttctggtgtccattgaaaatagccactggagaagtcttcaac
ccagtcatattaaacatgtagatgttatcgtctgtgcaattagcaaataaagtagagccagtggaatccaaaatcagacttgaatatccaagttttcgagtgctg
ctacctgggtacaggaaagacttggatgctatgggttcttgtcgataagcagtataattcttacgtaaatcccatactttgattatcccatccacagctcctgctga
gactaaggtattctcgtcttgaaagaggaccacagtaacactttgctggaaatccacagaaggagcaagtccttttgaattctgtttcttcttgggttttgaaggg
gtttgcttgtctgaggtattgtgagctccactgatttgattcacttgcctataaaacccatctttttttgttgcacctggtatcccagaccataatgttgccatctcttccac
ccgtacagaatacagctttctcaaacttagaaaaggcaactgacttgaggctgcattgatgacctttgcatgttccaatcagctcaccagcttttacgtcccaaa
atttggctgtttgatcacctgctgctgtaacaagtttaagttcaccaggaacccaggccaggtcaaagacggcattccagtgagccatccattctttgaagcact
tctttctgaaactttgtgattctgtgttatacaatcgaacaaagccttcttcattggcaactgctagtacatgttccatattgggagcagaagagaaggtacatcca
aaaggaggaactgggactcctgtttctccataagaagtgtgttcatcattaccactgcactgataaccagtcagaagggattgaagagggtattgtgaagac
catccatttctcaggacgccaagctggggctggcggagcaccgaattgaagagtgcatagtccgggacgtcatagggataaaccatggtggccacgtcgt
attaatttccacgtgccagtaagcagtgggttctctagttagccagaaggtacctttacgagggtaggaagtggtacggaaagttggtataagacaaaagtgt
tgtggaattgaagtttactcaaaaaatcagcactcttttataggcgccctggtttacataagcaaagcttatacgttctctatcactgatagggagtaaactggat
atacgttctctatcactgatagggagtaaactgtagatacgttctctatcactgatagggagtaaactggtcatacgttctctatcactgatagggagtaaactcc
ttatacgttctctatcactgatagggagtaaagtctgcatacgttctctatcactgatagggagtaaactcttcatacgttctctatcactgatagggagtaaactcg
aggtgataattccactcgagtggctccggtgcccgtcagtgggcagagcgcacatcgcccacagtccccgagaagttgggggggaggggtcggcaattga
accggtgcctagagaaggtggcgcggggtaaactgggaaagtgatgtcgtgtactggctccgcctttttcccgagggtgggggagaaccgtatataagtgc
agtagtcgccgtgaacgttctttttcgcaacgggtttgccgccagaacacaggtgtcgtgacgcgggatccgccaccatggattacaaagacgatgacgat

aagatgtctagactggacaagagcaaagtcataaactctgctctggaattactcaatggagtcggtatcgaaggcctgacgacaaggaaactcgctcaaa
agctgggagttgagcagcctaccctgtactggcacgtgaagaacaagcgggccctgctcgatgccctgccaatcgagatgctggacaggcatcataccc
actcctgccccctggaaggcgagtcatggcaagactttctgcggaacaacgccaagtcataccgctgtgctctcctctcacatcgcgacggggctaaagtg
catctcggcacccgcccaacagagaaacagtacgaaaccctggaaaatcagctcgcgttcctgtgtcagcaaggcttctccctggagaacgcactgtacg
ctctgtccgccgtgggccactttacactgggctgcgtattggaggaacaggagcatcaagtagcaaaagaggaaagagagacacctaccaccgattctat
gcccccacttctgaaacaagcaattgagctgttcgaccggcagggagccgaacctgccttccttttcggcctggaactaatcatatgtggcctggagaaaca
gctaaagtgcgaaagcggcgggccgaccgacgcccttgacgattttgacttagacat
